# MacaSurfer: unified surface-volume mapping of the macaque brain across the lifespan

**DOI:** 10.64898/2026.06.14.732101

**Authors:** Yahui Wei, Haiyan Wang, Yufan Wang, Lixin Chen, Luqi Cheng, Jinquan Gao, Qi Zhu, Congying Chu, Ting Xu, Chaohong Gao, Tianzi Jiang, Wim Vanduffel, Lingzhong Fan

## Abstract

Macaque brain MRI is central to translational and comparative neuroscience, yet multi-site, longitudinal, and cross-species analyses are hindered by a lack of unified, automated structural processing tools. Existing pipelines, mostly adapted from human neuroimaging or restricted to fragmented steps, fail to provide robust surface-volume representations across heterogeneous acquisitions and developmental stages. Here we introduce MacaSurfer, a fully automated, containerized framework for unified surface-volume mapping of the macaque brain across the lifespan. MacaSurfer features components tailored for macaque anatomy: a tissue segmentation model, a tissue-guided bias-field correction method optimizing structural mapping from T1-weighted images alone, topology-aware surface reconstruction, and surface-aware volumetric registration. Validated on 1,346 imaging sessions from 965 macaques across 39 international sites (spanning 2 weeks to 23 years of age), MacaSurfer demonstrated exceptional anatomical consistency, test-retest precision, and robustness against image degradation. Leveraging MacaSurfer-derived morphometry, we established normative trajectories from 835 macaques, providing a standardized reference for downstream individualized deviation analysis. MacaSurfer is openly available with source code, containers, and pretrained models, offering a reproducible ecosystem to accelerate developmental, translational, and comparative neuroimaging.

## Introduction

Macaques play an important role in translational and comparative neuroscience. Their phylogenetic proximity to humans, together with the feasibility of experimental manipulation, makes them particularly valuable for studying cortical organization, brain development, disease mechanisms and cross-species homology^1–3^. Within this framework, structural MRI provides an essential in vivo tool for comparing individuals, tracking structural change over time and relating macaque brain organization to human neuroimaging phenotypes^4^. Therefore, macaque structural MRI is not only a tool for describing anatomy, but also a framework for organizing downstream analyses across subjects, studies and species.

Despite this importance, macaque MRI datasets are typically acquired in relatively small cohorts and under site-specific experimental conditions^5^. Substantial variation in scanner hardware, acquisition protocols, spatial resolution, image quality and coil-related intensity inhomogeneity further complicates data processing and limits direct comparability across studies^6^. The scientific value of existing datasets, and their utility for future work, therefore depends not only on data acquisition itself, but also on whether diverse datasets can be processed with a unified and reliable workflow that generates anatomically consistent representations across studies^7^. Such a workflow is essential for enabling direct comparison, reproducibility and the cumulative reuse of macaque neuroimaging data.

However, this level of integration remains lacking in existing approaches, which are largely adapted from human MRI analysis or focused on isolated processing steps. While tools such as CIVET-Macaque^8^ and PREEMACS^9^ have advanced automated cortical surface extraction, and recent deep learning methods have improved brain extraction and tissue segmentation^10,11^, these developments do not yet provide a unified preprocessing workflow that can robustly handle cortical surface representations with volumetric anatomy across heterogeneous acquisitions, image qualities and developmental stages.

We therefore developed MacaSurfer, an end-to-end macaque-specific framework that integrates preprocessing, cortical and subcortical segmentation, topology-aware surface reconstruction, multi-atlas parcellation, subject-to-template mapping, morphometric and volumetric quantification, visual quality control and normative reference analysis within a single reproducible workflow. To address recurrent failures in macaque MRI, MacaSurfer combined deep-learning-based anatomical priors with geometry-aware reconstruction strategies, including tissue-guided bias-field correction for reliable structural mapping from T1-weighted images alone, topology-preserving signed-distance representations for robust surface modeling, and surface-aware volumetric registration to improve cortical correspondence while preserving volumetric alignment. We validated MacaSurfer on 1,346 imaging sessions from 965 rhesus macaques acquired at 39 international sites, spanning ages from 2 weeks to 23 years, and further established normative developmental trajectories from 835 macaques comprising 1,145 sessions across 26 sites. Together with its release of source code, containers, pretrained models and normative resources, MacaSurfer is intended not only as a practical processing workflow, but also as a shared infrastructure for integrating existing macaque MRI datasets and supporting downstream developmental, translational and comparative studies (https://github.com/yahuiwei123/MacaSurfer).

## Results

### MacaSurfer establishes a unified framework for macaque surface–volume mapping

We first sought to establish MacaSurfer as a unified framework for structural MRI analysis of the macaque brain, rather than as a tool for an isolated preprocessing step. To resolve the systemic methodological fragmentation in non-human primate brain imaging research, we built MacaSurfer as a standardized, fully automated and high-throughput computational workflow that spans preprocessing, cortical and subcortical segmentation, surface reconstruction, parcellation, subject-to-template mapping, morphometric quantification, visual quality control and downstream normative analysis within a single reproducible pipeline. By organizing cortical surface geometry and volumetric anatomy within a common representation, the framework supports anatomically consistent downstream analysis across subjects, studies and imaging conditions. Built upon a containerized Nextflow orchestration engine^12^, MacaSurfer achieves end-to-end automation from raw structural inputs to standardized morphometric outputs, supports process-level caching and parallel execution across local workstations and high-performance clusters, and is BIDS-compliant^13^ — automatically configuring its workflow based on available inputs while retaining the flexibility for targeted customization in atypical or challenging cases.

To maximize empirical robustness across heterogeneous datasets and complex edge cases, MacaSurfer integrates an automated quality control system coupled with specialized remediation protocols. The workflow records all processing parameters and generates comprehensive visual reports at every critical stage (brain masking, tissue segmentation and surface fitting), enabling rapid and transparent cohort assessment. When confronting common failure modes (e.g., skull-stripping errors, white-matter defects) or severe structural anomalies (e.g., ventricular enlargement, extreme artifacts), MacaSurfer’s modular architecture provides targeted remediation procedures; these individual modules can be iteratively refined and re-executed without recomputing the entire pipeline, drastically reducing the cost of manual intervention. The workflow supports GPU acceleration while remaining fully compatible with CPU-only environments, and the complete source code and comprehensive documentation are openly available to foster reproducible research and community-driven advancement.

### MacaSurfer generalizes across sites, ages and challenging anatomical conditions

To evaluate the generalizability and robustness of MacaSurfer, we applied it to a large multi-center collection comprising 1,346 imaging sessions from 965 rhesus macaques acquired at 39 international research sites. This cohort represents a wide spectrum of acquisition heterogeneities, demonstrating the workflow’s immunity to variations in scanner hardware, coil configuration and imaging protocol (Fig. 2a).

**Figure 1.**
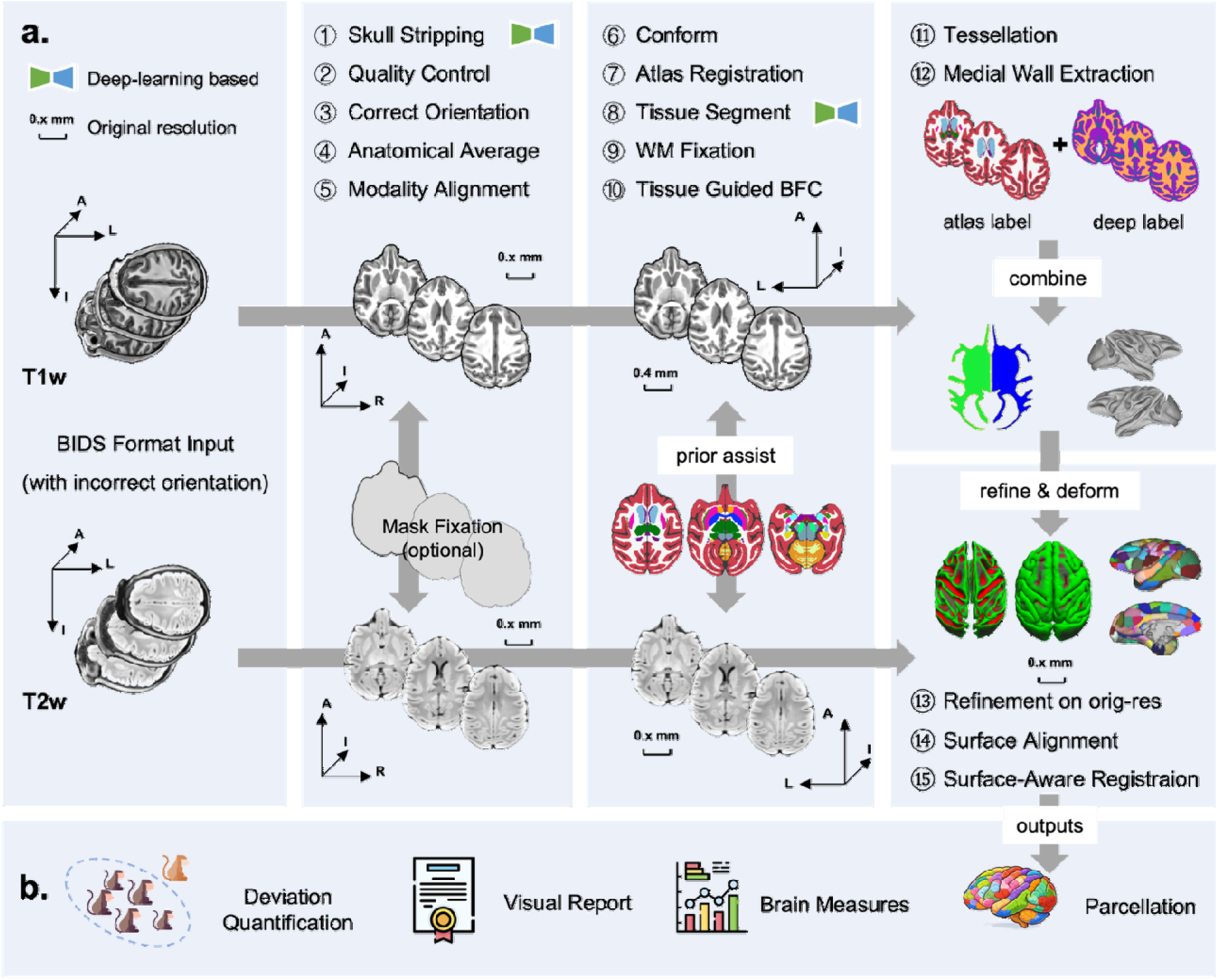
The MacaSurfer Workflow. (a) Pipeline Architecture. The workflow proceeds through four distinct, automated phases: Data Preparation (involving automated orientation correction, skull stripping, and within-subject rigid alignment); Segmentation and Enhancement (incorporating deep semantic priors for 18-class tissue segmentation, hybrid vessel detection, and tissue-guided bias field correction to resolve intensity inhomogeneities); Surface Initialization (performing template-space tessellation, SDF-based white matter skeletonization, and graph-based medial wall definition); and Surface Refinement (executing multi-contrast surface deformation and surface-aware volumetric registration to ensure precise sulcal alignment). **(b) Comprehensive Analytical Outputs.** MacaSurfer provides high-resolution cortical and subcortical parcellations across multiple coordinate spaces (native, ACPC, and template), supported by automated slice-wise visual quality control reports. The pipeline generates granular region-wise and vertex-wise morphometric measurements (e.g., thickness, area, volume, and sulcal depth). Crucially, it incorporates normative developmental trajectories derived from 835 individuals to generate subject-specific Z-score maps, enabling quantitative assessment of individual anatomical deviations relative to population baselines.

**Figure 2.**
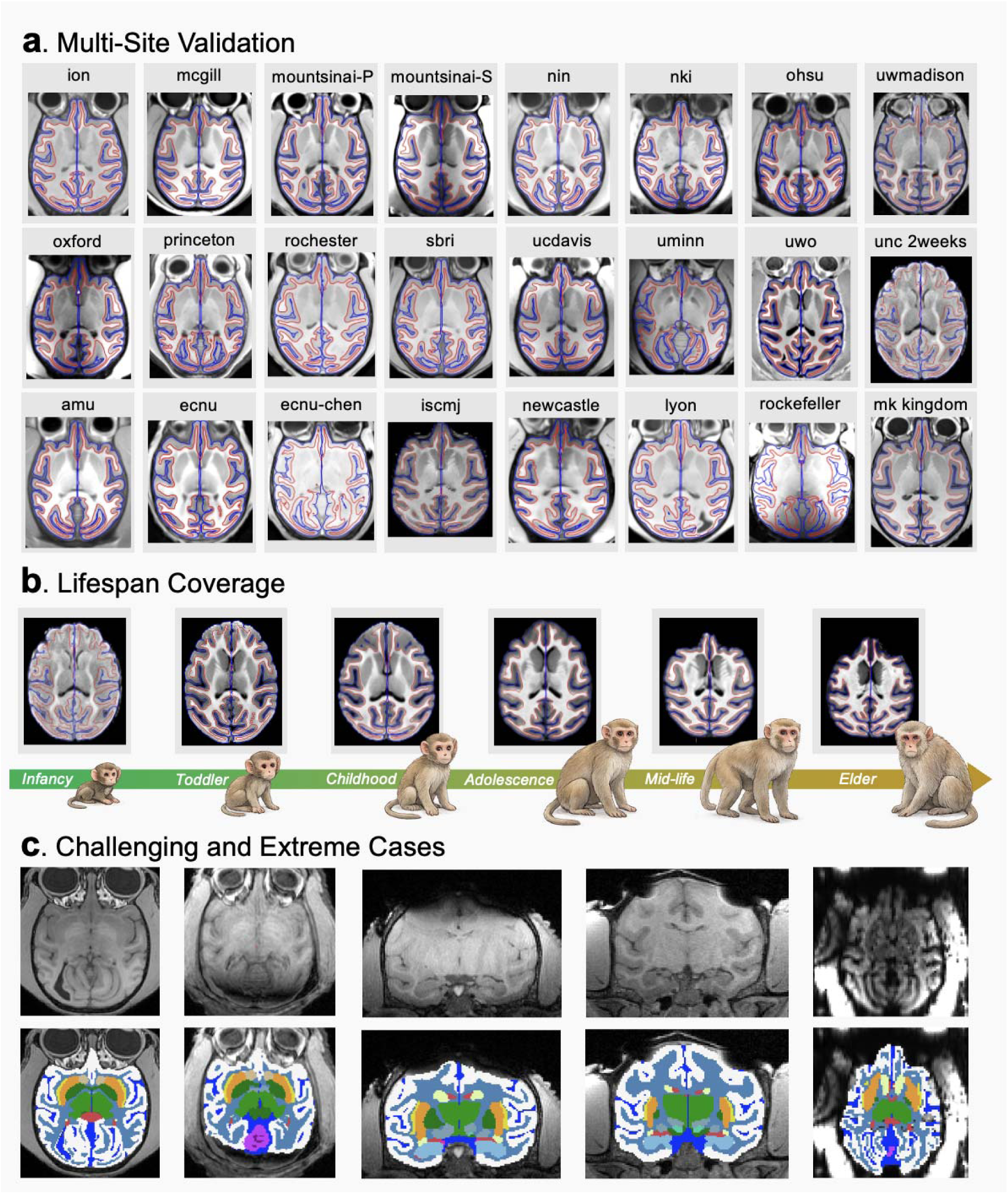
MacaSurfer’s Generalizability Across Acquisition Heterogeneity, Lifespan, and Edge Cases. **(a)** Representative axial T1-weighted slices with reconstructed white matter (red) and pial (blue) surfaces overlaid from 24 representative international sites, showcasing robustness against varying contrast and resolution across a total of 39 sites. **(b)** Reconstructions spanning the macaque lifespan—from 2-week-old infants to aged adults (up to 23 years)—demonstrate stable performance despite substantial age-related anatomical variations in cortical thickness and gyrification. **(c)** Examples of challenging “edge cases,” including images with significant artifacts, low signal-to-noise ratios, or atypical anatomy, where the pipeline successfully recovers plausible cortical and subcortical boundaries.

**Figure 3.**
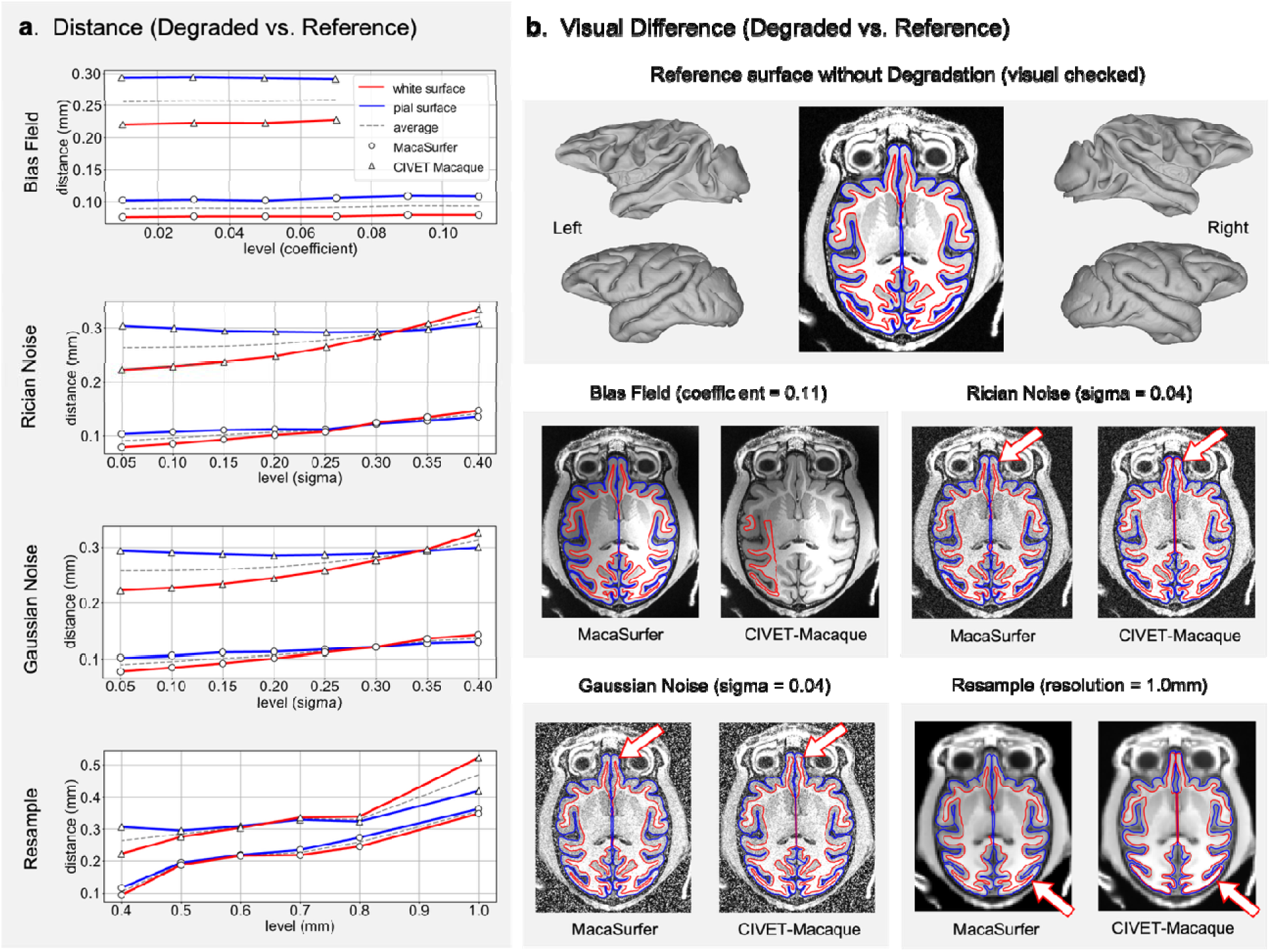
Quantitative robustness to image degradation. **(a)** Performance curves displaying the symmetric surface-to-surface distance (error) between reconstructed surfaces and the ground truth under increasing levels of image degradation (bias field, noise, and downsampling). MacaSurfer (circles) consistently exhibits lower error rates compared to CIVET-Macaque (triangles) for both white matter (red) and pial (blue) surfaces. **(b)** Visual comparison of surface reconstructions under noisy conditions. Red arrows indicate specific region where the comparison method (CIVET-Macaque) fails to capture correct cortical topology or suffers from boundary erosion due to N4 correction limitations, whereas MacaSurfer’s Tissue-Guided Correction maintains high boundary fidelity.

As illustrated by the representative reconstructions from 24 sites, MacaSurfer produced anatomically consistent cortical surfaces despite substantial heterogeneity in image contrast and resolution. The workflow further exhibited remarkable stability across the macaque lifespan: high-fidelity reconstructions were successfully obtained from 2-week-old infants — whose brain images typically feature low tissue contrast due to incomplete myelination — through to aged adults (23 years; Fig. 2b). The framework’s ability to maintain reconstruction fidelity in early postnatal brains is particularly notable, as infant macaque MRI presents unique challenges (reduced gray–white matter contrast, higher water content and smaller brain volume) that frequently lead to under-segmentation or topological defects in conventional pipelines.

We then asked whether MacaSurfer could handle challenging “edge cases” in which reliable cortical surface reconstruction may be intrinsically difficult or inappropriate, because distorted anatomy or degraded image quality can compromise tissue boundaries and surface topology. We therefore evaluated whether the framework could still provide anatomically meaningful structural outputs in scans exhibiting atypical ventricular enlargement, scan-related anatomical deformation or severe motion artifacts. The macaque-specific tissue segmentation module produced plausible cortical, subcortical and ventricular labels in these challenging scans, enabling reliable identification of major anatomical compartments even when downstream surface reconstruction was limited or required cautious interpretation (Fig. 2c). In subjects with enlarged ventricles, the segmentation preserved the separation between ventricular space, periventricular tissue and cortical gray matter; in scans affected by deformation or motion, the model still recovered the main tissue compartments despite local signal degradation. These results indicate that MacaSurfer provides robust structural segmentation for heterogeneous macaque MRI datasets and can support volumetric analysis in cases that may not be suitable for standard cortical surface reconstruction.

### Macaque-tailored components improve structural mapping and anatomical correspondence

Then, we evaluated whether the macaque-specific components of MacaSurfer improved structural mapping at five key failure points: registration robustness under incorrect anatomical orientation; systematic benchmarking and selection of registration tools for cross-modal alignment; tissue segmentation accuracy for downstream processing; bias-field correction in low-contrast regions; and topology-preserving surface reconstruction in compact, highly folded cortices.

### Automatic orientation correction eliminates registration failures caused by mislabeled anatomical coordinates

In 39 scans from six data collections with randomly permuted anatomical headers (Methods), our correction algorithm recovered the correct orientation in all cases (100%). After correction, affine registration to the MEBRAIN T1w template yielded consistently higher normalized cross□Jcorrelation (NCC; mean = 0.54 ± 0.09) than without correction (mean = 0.21 ± 0.13; 32/39 sites showed unacceptably low NCC or failed registration). These results demonstrate that automatic orientation correction markedly reduces inter□Jsite registration variability (Supplementary Fig. 4 and 5).

### Systematic benchmarking of registration tools ensures robust cross-modal alignment

We benchmarked three widely used registration tools across seven parameter configurations for intra□J and cross-modal alignment (Supplementary Table 1). While all tools performed comparably for T1w□Jto□JT1w and T2w□Jto□JT2w registration (Supplementary Fig. 6a, b), cross□Jmodal T2w□Jto□JT1w registration revealed substantial differences: the highest visual pass rate was achieved by NiftyReg reg_aladin (100%), followed by ANTs (97.0%) and FLIRT (93.1%) (Supplementary Fig. 6c). Based on these results, we adopted FLIRT for intra□Jmodal rigid alignment and contrast□Jenhanced reg□aladin for cross□Jmodal T2w□Jto□JT1w registration in the MacaSurfer workflow.

### Deep-learning-based tissue segmentation provides robust priors and initial topology

In within-site cross-validation and cross-site generalization experiments, the trained model achieved high Dice similarity coefficients, confirming that the segmentation model generalizes across acquisition protocols and field strengths. This robust segmentation stabilized all downstream steps.

### Tissue-guided bias-field correction recovers white matter in low-contrast regions

Compared with standard N4 correction, MacaSurfer’s tissue□Jguided bias□Jfield modeling recovered high□Jfrequency intensity variations within deep cortical folds and prevented boundary erosion in the occipital lobes, preserving thin white matter strands that conventional correction often removed (Supplementary Fig. 7). This substantially improved white□Jsurface placement in regions where standard correction degrades surface initialization.

### Topology-preserving surface modeling recovers thin and elongated white matter and yields anatomically correct medial walls

By enforcing a continuous white matter skeleton via signed distance constraints, MacaSurfer recovered thin, elongated white matter in the occipital and anterior frontal lobes that voxel□Jwise segmentation frequently misses (Supplementary Fig. 8). For the medial wall, graph□Jbased extraction of the largest connected component produced anatomically correct cortical masks: unlike standard FreeSurfer reconstructions, MacaSurfer consistently excluded medial wall structures, yielding a cortical boundary that strictly adheres to anatomical limits (Supplementary Fig. 9).

### Surface-aware volumetric registration improves cross-subject parcel alignment

We finally tested whether the reconstruction gains produced by our surface-aware volumetric registration algorithm translated to improved cross-subject anatomical correspondence. Volume-based registration and smoothing can degrade cortical spatial localization when neighboring cortical areas are close in Euclidean space but separated along the folded sheet; however, purely surface-based processing is often insufficient for macaque MRI because functional, diffusion and subcortical measurements are acquired volumetrically, and volume-to-surface projection introduces sampling bias. To address this, MacaSurfer’s symmetric diffeomorphic registration was augmented with a surface-aware loss that jointly constrains the deformation field to align cortical surfaces while maintaining volumetric smoothness. Warping native Macaque Brainnetome Atlas^14^ (MacBNA) 124 parcellations to template space in 12 subjects from the Monkey Kingdom dataset, the surface-aware approach yielded a significantly higher mean Dice coefficient across 124 regions (76.02□J%□J±□J9.27□J%) than volume-only registration (70.31□J%□J±□J11.20□J%) (Supplementary Fig.□J10), confirming that incorporating cortical geometry into volumetric deformation yields more accurate individual-to-template mapping.

### MacaSurfer is robust under image degradation and reproducible across repeated scans

To quantify robustness and reproducibility, we evaluated MacaSurfer under three complementary conditions: controlled image degradation, repeated acquisitions from the same individuals, and a T1w-only versus combined T1w/T2w comparison.

### Robustness under controlled image degradation

To quantify the stability of MacaSurfer under suboptimal imaging conditions, we performed a stress test using 12 repeated scans of a single subject from the Monkey Kingdom dataset. We generated degraded T1-weighted variants via controlled perturbations — bias-field distortion, Gaussian and Rician noise, and spatial downsampling — and compared MacaSurfer’s performance with CIVET-Macaque^8^ without prior denoising, to ensure a stringent evaluation. MacaSurfer consistently yielded lower symmetric surface-to-surface distance errors across all noise types and severity levels than the traditional pipeline (Fig. 4). This superior performance is primarily attributable to the integration of semantic tissue priors and the tissue-guided bias-field correction, which allow MacaSurfer to disentangle coil-induced inhomogeneities from anatomical signals and preserve topological stability even in low-contrast temporal and occipital regions, where standard intensity-based approaches typically fail.

**Figure 4.**
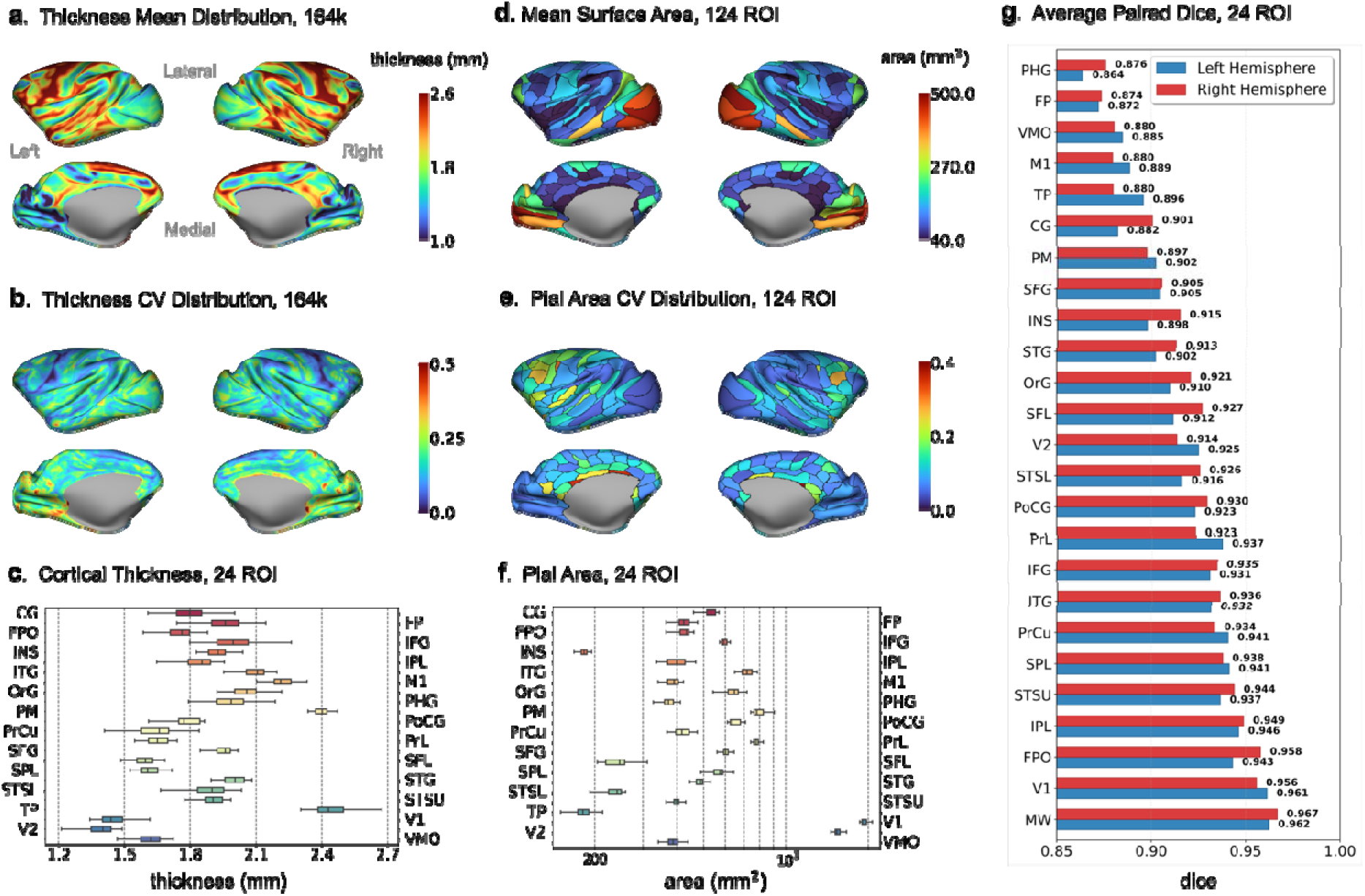
Test-retest reliability of cortical metrics. The precision of the workflow using repeated scans from the Site-NIN dataset, with all results displayed on the fs_LR template. **(a)** Mean cortical thickness maps averaged across repeated scans. **(b)** Vertex-wise coefficient of variation (CV) for cortical thickness, highlighting areas of high spatial stability. **(c)** Boxplots showing the distribution of cortical thickness across 24 standard regions of interest (ROIs). **(d-e)** Mean pial surface area and its corresponding CV for 124 brain regions. **(f)** Log-scale distribution representing the variability of surface area across different ROIs. **(g)** Mean pairwise Dice coefficients for 24 standard ROIs, confirming the high spatial overlap and reproducibility of volumetric segmentations across independent scan sessions.

### Reproducibility across repeated scans

Reliability in longitudinal and translational studies requires that surface metrics remain stable across repeated measurements. We evaluated the intrasubject reproducibility of MacaSurfer using the Site-NIN dataset, analyzing two rhesus macaques with 24 and 42 repeated scans, respectively, at 0.6 mm isotropic resolution (matching the physical voxel size of the scans). Each scan was reconstructed independently in its native space without inter-scan averaging. Under these conservative conditions, MacaSurfer achieved submillimeter precision: regional cortical thickness deviations typically remained below 0.6 mm, and we observed consistently low coefficients of variation (CV) for cortical thickness and high Dice similarity coefficients for volumetric segmentations across repeated acquisitions (Fig. 4). This precision supports the use of MacaSurfer for longitudinal monitoring — for example, tracking subtle neurodegenerative progression — and provides a stable scaffold for individualized normative modelling.

### Applicability to T1w-only acquisitions

We further asked whether MacaSurfer could maintain high-fidelity structural mapping from T1-weighted images alone. We compared surface reconstructions derived from T1w-only inputs against those obtained from combined T1w/T2w inputs in 80 subjects across 12 independent imaging sites. Vertex-wise paired t-tests (FDR-corrected, p < 0.01) revealed no significant differences in mean cortical thickness across 97.12% and 97.79% of left and right hemisphere cortical vertices, respectively, and complementary Kolmogorov–Smirnov tests showed that cortical thickness distributions were statistically indistinguishable across more than 99.9% of the cortex in both hemispheres. Other metrics (e.g., sulcal depth) exhibited slightly higher rates of significant difference (∼11–13%), but cortical curvature and thickness remained remarkably consistent across modalities (Supplementary Fig. 11; Supplementary Result 3). This parity supports the use of MacaSurfer in datasets where multimodal structural acquisitions are unavailable or inconsistent, including historical archives and simplified acquisition protocols.

### Normative Developmental Trajectories Enable Individualized Anatomical Assessment

To alleviate the computational and manual inspection burden that large-scale macaque data processing would otherwise impose on individual studies, and to provide a standardized downstream reference for MacaSurfer-based analyses, we finally constructed normative developmental trajectories from structurally processed macaque MRI data. Using MacaSurfer-derived morphometric features from a multi-site cohort of 835 neurologically healthy macaques, comprising 1,145 imaging sessions across 26 international imaging sites, we trained 2,942 regional normative models covering cortical and subcortical structures across both hemispheres and four parcellation schemes (M129, M132, MacBNA124, the multi-modal combined atlas, and subcortical aseg labels). Model performance was evaluated by cross-validated out-of-sample estimates on a held-out test set, using explained variance (EXPV), mean absolute error (MAE), root mean squared error (RMSE), coefficient of determination (R²) and Spearman’s correlation.

Model accuracy varied systematically across morphometric modalities (Supplementary Table 3). Among cortical features, gray matter volume and cortical thickness showed the strongest predictive performance (mean EXPV 0.25 ± 0.09 and 0.24 ± 0.10, respectively), and gray matter area showed comparable performance (mean EXPV 0.22 ± 0.07). Sulcal depth and cortical curvature exhibited lower predictive accuracy (mean EXPV 0.11 ± 0.06 and 0.06 ± 0.06), consistent with greater inter-individual variability and weaker age-related signal. Subcortical volume models achieved the highest overall performance, with a mean EXPV of 0.39 ± 0.12 and a mean Spearman’s ρ of 0.58 ± 0.11.

Model performance also showed substantial regional heterogeneity (Supplementary Table 4). For cortical thickness models in the MacBNA124 atlas, several temporal, cingulate, orbital, precuneus and prefrontal regions showed moderate-to-high predictive accuracy, including the inferior temporal gyrus (EXPV = 0.52, ρ = 0.57), ventral rostral cingulate gyrus (EXPV = 0.45, ρ = 0.56), rostromedial orbital gyrus (EXPV = 0.43, ρ = 0.57), dorsal intermediate precuneus (EXPV = 0.42, ρ = 0.58) and ventromedial prefrontal cortex (EXPV = 0.42, ρ = 0.59). Lower performance was observed in regions such as the ventral rostral temporal pole, ventral rostral insular cortex and lateral rostral temporal pole, where EXPV ranged from 0.02 to 0.04. Performance was broadly consistent across cortical parcellation schemes, with average EXPV ranging from 0.25 ± 0.19 (MacBNA124) to 0.28 ± 0.22 (M129); the multi-modal combined atlas achieved the highest average performance (EXPV 0.41 ± 0.32), driven primarily by improved performance in surface area and cortical thickness models.

Overall, 62% of regional models attained an explained variance greater than 0.4, providing a robust reference against which individual animals can be evaluated. By generating subject-specific Z-score maps that quantify deviations from age-informed expected trajectories, this normative framework extends the utility of MacaSurfer beyond structural preprocessing alone, supporting individualized assessment, longitudinal monitoring and the analysis of intervention-related structural change in non-human primate (NHP) models. Detailed evaluation and selection of the normative models, including metric-specific, regional and atlas-specific performance and hyperparameter settings, are provided in Supplementary Result 1. Together, these results show that MacaSurfer provides not only robust structural processing across heterogeneous macaque MRI datasets, but also a standardized anatomical framework for downstream developmental, translational and comparative studies of the macaque brain.

## Discussion

This study establishes MacaSurfer as a comprehensive computational platform that addresses the long-standing methodological fragmentation in macaque neuroimaging^5,15^ by unifying surface and volume processing within a single, bidirectional information flow. Unlike traditional pipelines adapted from human-centric architectures ^16^ that treat volumetric segmentation and surface reconstruction as independent, feedforward stages, MacaSurfer implements an integrated framework where volumetric deep-learning priors stabilize surface initialization, while cortical geometric constraints concurrently refine volumetric registration. Built upon a containerized Nextflow orchestration engine^12^, the framework achieves end-to-end automation from raw structural inputs to standardized morphometric outputs. This architectural convergence provides the robustness required to handle the high heterogeneity of multi-site NHP datasets across the entire lifespan^5^, from 2-week-old infants to aged adults, while process-level caching ensures computational efficiency.

A primary contribution of this work is the “expert-in-the-loop” bootstrapping paradigm, which offers a systematic solution to the chronic scarcity of ground-truth annotations in NHP research. We addressed the “circular dependency”—where robust tools require extensive labels, yet labels require reliable tools^17^—by using macaque-tailored algorithms to generate high-quality pseudo-labels. Through three rounds of expert neuroanatomical review and model retraining involving 2,157 scans from 39 international sites, we progressively refined a Swin UNETR-based deep learning model for 18-class tissue segmentation^18,19^. The model was initialized with weights from the VoCo (Volume Contrastive) self-supervised pre-trained model^29^, providing it with strong generalizability to anatomical variations and imaging artifacts. This approach would serve as a extensible framework for imaging other species with limited labels, providing a scalable roadmap for extending high-fidelity anatomical modeling across the primate lineage^20,21^.

This achievement is made possible by two design principles. The first is to systematically resolve species-specific acquisition and anatomical challenges that would otherwise corrupt the volumetric foundation. Automated orientation correction uses a two-stage FLIRT search to resolve coordinate header errors caused by non-supine scanning, achieving a 100% success rate across 39 sites. Systematic cross-modal rigid registration, for which benchmarking led to the selection of NiftyReg reg_aladin, directly overcomes the low tissue contrast typical of macaque MRI. We defined cortical and subcortical parcellations on the MEBRAINS template and, based on these, trained a dedicated deep learning segmentation algorithm tailored for macaques, thereby supplying robust anatomical priors for all subsequent processing. In human imaging, extreme inter-individual gyral variability forces a departure from Euclidean space via spherical intermediate mappings to align cortical geometry; the relative folding consistency in macaques instead permits simultaneous surface-volume alignment within a single Euclidean step^22–24^.

Therefore, the second principle exploits a defining macaque anatomical characteristic, namely the highly stereotyped cortical folding^25^, to achieve a deep, bidirectional integration of volumetric and surface-based optimization. This same regularity underpins an interleaved closed-loop architecture in which volume and surface continuously refine each other. Volume-domain operations, including deep-learning tissue segmentation, tissue-guided bias field correction, and the mapping of volumetrically extracted medial walls onto the surface, directly stabilize surface placement and spherical registration. Surface-derived constraints feed back to the volume: topology-preserving white-matter skeletonization projects surface geometry to recover thin white-matter strands missed by voxel-wise segmentation, and the surface-aware registration itself prevents blurring across gyral banks that are adjacent in Euclidean space but distant along the cortical sheet. The refined volumes become improved substrates for subsequent surface fitting, establishing iterative cycles in which surface-informed volume enhancement reciprocally elevates surface quality, interlocking with the direct volume-driven surface improvements described above. This tight integration, grounded in macaque-specific regularity, closes the loop and renders the pipeline resistant to the error propagation typical of strictly feedforward architectures.

MacaSurfer’s modality-invariant reconstruction serves as a critical mechanism for leveraging the scientific value of legacy datasets and historical archives. While multimodal T1w/T2w acquisitions are ideal, many large-scale collections lack matched T2-weighted scans. By integrating tissue-guided bias-field correction—which adaptively selects between an adaptive scatter RBF algorithm for low-frequency bias and a tissue-guided GMM algorithm for high-frequency artifacts—and topology-preserving signed-distance functions, MacaSurfer produces T1-only reconstructions that are statistically indistinguishable from multimodal results across more than 97% of the cortex. The SDF-based white matter skeletonization is particularly effective at recovering thin white matter strands in the occipital and anterior frontal lobes that voxel-wise segmentation often misses due to partial volume effects.

Inspired by the GAMLSS-based brain chart framework recently established for the rhesus macaque lifespan^26^, we constructed normative developmental trajectories from 835 healthy macaques using our MacaSurfer-derived morphometry to provide a foundational scaffold for individualized preclinical assessment. By providing a standardized reference system—analogous to pediatric growth curves—this framework trained 2,942 regional normative models that enable researchers to generate subject-specific Z-score maps to quantify anatomical deviations. The models incorporated sex and species as fixed-effect covariates and utilized ComBat-GAM^27^ for multi-site harmonization, capturing biologically plausible age-related trajectories from birth to early adulthood. This shifts the utility of the pipeline from cross-sectional group-level morphometry to individualized quantitative tracking of disease progression, characterizing neurodevelopmental milestones, and evaluating experimental interventions with high quantitative precision.

Despite these advancements, several limitations warrant consideration. While optimized for rhesus macaques, extending the framework to other NHP species will require species-specific templates, atlases, and cross-species transfer learning. Additionally, while the unified representation provides a robust scaffold, future work must integrate multimodal data—specifically the fusion of diffusion and functional MRI within this surface-centric coordinate space—for a holistic understanding of the primate connectome. Furthermore, extreme anatomical malformations or ultra-high-resolution acquisitions remain underrepresented in the current training sets. Finally, as metadata becomes more comprehensive, incorporating additional covariates such as genetic background, diet, rearing conditions, and anesthesia protocols into normative models will further enhance their predictive precision.

In conclusion, MacaSurfer mitigates the reproducibility challenges associated with lab-specific scripts by adopting BIDS-compliant^13^, containerized, and transparent workflows. Through its “glass-box” design, which provides visual quality-control reports for every stage (brain masking, tissue segmentation, and surface fitting), the platform ensures that both intermediate and final outputs are accessible for verification. By unifying surface and volume processing and leveraging iterative expert refinement, MacaSurfer establishes the necessary computational infrastructure to accelerate the discovery of conserved neurobiological principles across the primate lineage. This standardized ecosystem ultimately strengthens the translational bridge to human neuroscience, facilitating large-scale, cross-species brain mapping.

## Methods

### Framework architecture and design

MacaSurfer is a modular, configuration-driven pipeline for macaque cortical reconstruction that supports both multimodal (T1w + T2w) and unimodal (T1w-only) inputs (Fig. 5b). The workflow is organized around four integrated stages (Fig. 1a). **(1) Data Preparation:** an automated orientation correction module resolves coordinate header errors prevalent in non-supine macaque acquisitions by registration-based alignment to the MEBRAINS template, followed by within-subject rigid alignment across repeated scans. **(2) Quality Enhancement:** a 3D deep-learning engine jointly segments 18 tissue classes; these semantic priors drive a Tissue-Guided Bias Field Correction algorithm that decouples coil-induced intensity inhomogeneities from biological signals, recovering white-matter contrast in low-signal temporal and occipital regions. **(3) Surface Reconstruction:** signed distance functions (SDFs) and template-derived priors constrain a topologically correct white-matter skeleton, recovering thin white-matter strands frequently lost to partial volume effects in voxel-wise segmentation. Following white-surface reconstruction, volume-to-surface mapping combined with vertex connectivity is used to extract middle-wall voxels, after which the pial surface is reconstructed—refined on T2w data when available. **(4) Spatial Transformation:** Surface-Aware Volumetric Registration constrains 3D deformation fields with explicit cortical geometric correspondences, improving cross-subject alignment in complex sulcal regions compared to conventional intensity-based registration. Surface alignment is achieved via MSM spherical registration to the template surface. The detailed workflow is summarized in Fig.1a, and algorithmic components are described in subsequent sections.

A central design principle of MacaSurfer is the establishment of a unified anatomical reference that links template-space atlas definitions, subject-level label generation, and population-level normative modeling within a single consistent framework. We constructed this reference in MEBRAINS template space^28^, selected for its high-quality in vivo T1w/T2w contrast. A hybrid atlas was built by combining subcortical annotations—derived from the MEBRAINS gray–white matter boundary and augmented with nuclei definitions transferred from the NMT-based Subcortical Atlas of the Rhesus Macaque (SARM) ^29–31^ via symmetric diffeomorphic registration^22^ followed by manual refinement—with cortical parcellations based on the Macaque Brainnetome Atlas^14^, transferred from CIVM (Center for In Vivo Microscopy) space^33^ to the MEBRAINS spherical surface and projected onto the MEBRAINS mesh. This produced a unified reference in which cortical labels are surface-consistent and subcortical labels are volumetrically defined, ensuring that the same anatomical definitions drive both individual processing outputs and population-level morphometric measurements.

To overcome the fundamental scarcity of high-quality annotated macaque MRI data, we developed an iterative, closed-loop label-generation and model-refinement framework (Fig. 5). Structural MRI data were aggregated from public and private sources—including PRIME-DE^5^, the UNC–Wisconsin developmental cohort^34^, OpenNeuro^35^, and private acquisition sites—spanning broad variation in age, modality, image quality, and acquisition protocol. Images were first processed with an initial MacaSurfer workflow to generate volumetric and surface-derived annotations. Expert neuroanatomists then reviewed these outputs to identify recurrent failure modes: incomplete skull stripping, tissue underestimation, inaccurate tissue boundaries, and topological defects affecting surface reconstruction. Based on these findings, processing parameters and strategies were refined, and the workflow was re-run. Across three rounds of reconstruction, expert review, and parameter refinement, this process generated progressively improved, anatomically consistent labels optimized for surface reconstruction.

The final curated image–label pairs were used to train macaque-specific deep learning models for skull stripping and tissue segmentation, initialized from a VoCo-pretrained backbone^36,37^ and trained with extensive spatial and intensity augmentations to ensure robustness across heterogeneous acquisitions (Data augmentation strategy, label generation method and training procedures are detailed in Fig. 5e, 5c, and 5d, respectively). After validation, the updated models were reintegrated into MacaSurfer, replacing the previous generation of priors and thereby closing the iterative loop. This closed-loop architecture—in which improved labels train better models, and better models produce improved labels—enables MacaSurfer to improve continuously as additional data and annotations become available. Notably, the overall pipeline architecture and processing logic remained stable across all three development rounds: subsequent iterations focused on refining targeted solutions to specific failure modes rather than restructuring the entire workflow, reflecting the initial design’s grounding in the most prevalent challenges of macaque structural MRI processing. The inference code and pretrained weights for all macaque-specific models are publicly released and can be used either within MacaSurfer or as standalone tools for structural MRI preprocessing.

**Figure 5.**
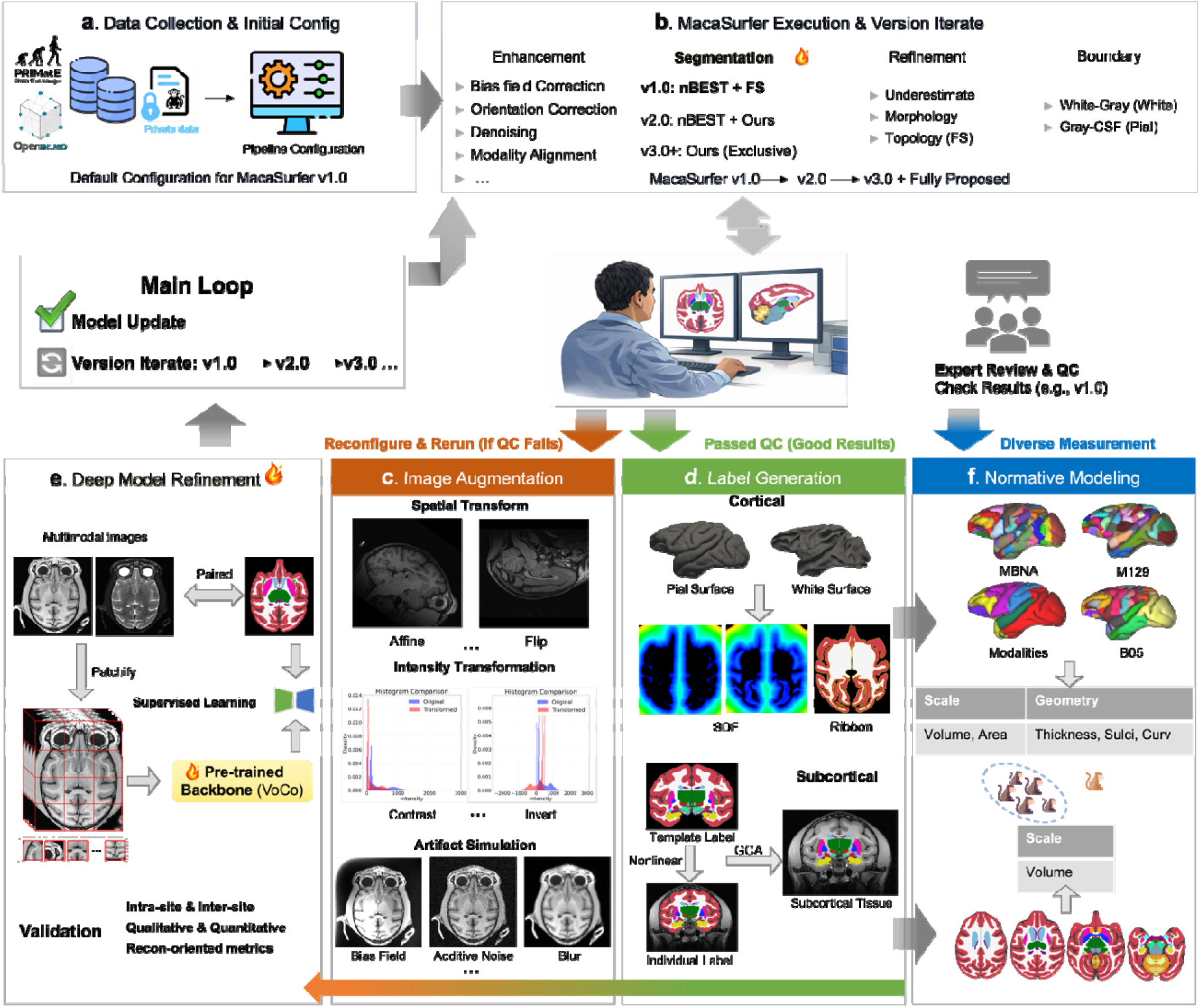
Closed-loop label generation framework and deep model refinement strategy in MacaSurfer. This schematic outlines the iterative bootstrapping paradigm designed to overcome the annotation scarcity bottleneck in non-human primate neuroimaging. By integrating an expert-in-the-loop feedback mechanism, the framework progressively refines initial segmentations into high-fidelity ground-truth labels, which subsequently drive the continuous evolution of domain-adapted deep learning models. **(a) Data Aggregation:** Raw structural MRI scans are pooled from highly heterogeneous public repositories and private clinical archives, establishing a diverse multi-site data foundation. **(b) Pseudo-label Generation:** Initial pipeline iterations (e.g., v1) perform baseline image enhancement, tissue classification, and surface reconstruction to generate preliminary, anatomically consistent “pseudo-labels” for cortical and subcortical structures. **(c) Human-in-the-Loop Refinement:** Expert neuroanatomists manually review the generated pseudo-labels and guide the application of automated topological correction procedures, such as fixation. Combined with dataset-specific parameter configurations, this rigorous curation converts preliminary outputs into high-fidelity ground-truth labels. **(d) Deep Model Training & Augmentation:** The curated labels, paired with heavily augmented raw data (employing domain randomization techniques), are utilized to supervise and fine-tune macaque-specific deep neural networks (e.g., skull-stripping and tissue segmentation models). **(e) Pipeline Integration & Evolution:** The validated, high-performance deep learning models are injected back into the core MacaSurfer engine, directly driving the iterative upgrade of the processing pipeline (e.g., to v2, v3) for exponential accuracy improvements. **(f) Final Outputs & Application:** The mature closed-loop framework ultimately produces highly accurate, generalized cortical and subcortical parcellations. These robust representations directly empower large-scale, individualized normative modeling and precise anatomical quantification across diverse acquisition conditions.

### MacaSurfer Technical Details

Although MacaSurfer underwent three rounds of iterative development, its overall architecture and processing logic remained largely unchanged. This was because, from the first version, the pipeline was designed around the most common and challenging problems in macaque structural MRI processing. Subsequent iterations therefore focused on refining targeted solutions to part of problems, rather than restructuring the entire workflow.

#### Preprocessing and Volumetric Analysis

Built on the unified framework architecture, we first perform standardized preprocessing and volumetric analysis to ensure reliable input for subsequent surface reconstruction. This section describes the key steps that correct spatial inconsistencies, extract brain tissues, and refine volumetric signals.

#### Automated Orientation Correction

To address the high frequency of coordinate header errors in macaque datasets^38^, each skull-stripped brain was rigidly registered to the MEBRAINS template^28^ using FLIRT^39–41^ to infer the native image orientation, which was then written to the input volume. This procedure was validated across all PRIME-DE acquisition sites^5^ by manual inspection. The detailed algorithmic workflow and registration settings are provided in Supplementary Note 1.

#### Volumetric Registration

Volumetric registration in MacaSurfer comprises three steps: (1) intra-subject rigid registration across repeated runs of the same modality; (2) rigid inter-modal registration between T1w and T2w images (when available); and (3) nonlinear registration of the individual T1w image to template space for atlas mapping. Based on systematic benchmarking across FSL’s FLIRT^39–41^, ANTs^32,42,43^, and NiftyReg^44^ (Supplementary Note 2), FLIRT was selected for intra-modality rigid registration and NiftyReg reg_aladin for cross-modality rigid alignment. Nonlinear template registration was performed using antsRegistration^32^, with FireANTs^45^ as an optional GPU-accelerated alternative for improved computational efficiency.

#### Unified Tissue Segmentation and Brain Masking

Accurate isolation and classification of brain tissues are critical for cortical surface reconstruction. MacaSurfer employs a unified multi-class tissue segmentation strategy in which brain extraction is implicitly defined by the union of predicted anatomical labels. The default model jointly segments cortical gray matter, white matter, cerebrospinal fluid, cerebellum, brainstem, and multiple subcortical structures, generating both a high-fidelity brain mask and anatomically informative labels for downstream analyses.

To address the scarcity of annotated macaque MRI data, tissue segmentation was developed using an iterative, pipeline-preserving bootstrapping strategy. The downstream reconstruction workflow (bias-field correction, volumetric registration, surface initialization, and surface refinement) was kept fixed while only the segmentation component was iteratively improved. Initial labels were obtained by combining deep learning–based cortical tissue segmentation^10^ with atlas-informed subcortical labeling after volumetric registration. Cases with suboptimal masks or tissue labels were manually corrected and reprocessed, yielding a curated set of anatomically consistent, surface-informed labels for model training.

Using this strategy, we assembled a training set of 2,157 macaque MRI scans from 39 acquisition centers. The final segmentation model is based on a SwinUNETR-B architecture^18,19^ initialized with self-supervised pretraining and fine-tuned for 18-class tissue segmentation. Complete details of the training dataset, model architecture, augmentation strategies, and performance metrics are provided in Supplementary Note 3.

### Optional Hybrid Segmentation Mode

An optional hybrid mode, in which cortical tissues are segmented via deep learning and subcortical structures are assigned by atlas-based probabilistic classification after volumetric registration, is available for compatibility with legacy workflows and method comparison (see Supplementary Note 3).

#### Surface Reconstruction and Refinement

Based on the high-quality volumetric outputs from preprocessing, we proceed to reconstruct and refine the cortical surfaces with strict topological constraints. This section presents the core algorithms that generate accurate white and pial surfaces while addressing macaque-specific anatomical challenges.

#### White Surface Initialization and Topology Correction

To generate the initial white surface, we construct a volumetric mask that provides a closed and topologically valid boundary for surface initialization. This mask is derived from the aseg volume produced by the hybrid tissue segmentation strategy, after all images have been conformed to the MEBRAINS template space^28^. By explicitly incorporating subcortical structures into the initialization mask, the resulting white surface maintains a globally closed topology suitable for subsequent deformation and refinement.

This volumetric representation ensures anatomically consistent boundaries at the cortex–subcortex interface and prevents topological defects that can arise when white matter is considered in isolation. Thin and elongated white matter structures—particularly in the occipital lobe and anterior frontal lobe—may be underrepresented in voxel-wise segmentation due to partial volume effects^46^. To recover these regions, we integrate template-derived white matter priors via a Signed Distance Function (SDF)^47^ computed from the template white matter surface projected into subject space. This skeleton acts as a hard topological constraint, enabling recovery of white matter strands missed by voxel-wise segmentation.

For hemisphere separation, we replaced the conventional sagittal cutting-plane strategy with a subject-specific hemispheric mask generated using FreeSurfer’s make_hemi_mask, which estimates a symmetric half-way space via inverse-consistent registration^48^ to avoid inaccuracies from planar midsagittal approximations^49,50^.

#### Tissue-guided Bias Field Correction

FreeSurfer’s white surface reconstruction relies heavily on intensity-based energy terms^51^, and residual bias fields after standard N4 correction^52,53^ vary in spatial scale across datasets. To address this heterogeneity, we introduced a Tissue-Guided Bias Field Correction framework that adaptively selects between two complementary strategies: high-frequency bias is modeled using a Gaussian Mixture Model (GMM) constrained by tissue segmentation labels, whereas low-frequency bias is estimated using radial basis function (RBF) mirror basis functions. Full algorithmic details and selection guidance are provided in Supplementary Note 4.

### Topological Correction and Medial Wall

Inaccuracies in volume-based atlas-to-subject nonlinear registration can cause the initial cortical surface mask (i.e., lh.cortex.label) to erroneously extend into the medial wall. Such misclassification between cortex and medial wall regions can adversely affect subsequent surface-to-surface registration, ultimately compromising the accuracy of mapping cortical parcellations onto individual subject surfaces^54^. To ensure an accurate and robust medial wall definition, we generate a precise midline mask by projecting the nonlinear hemisphere separation boundary (from volume space) onto the white surface. We then treat the surface as a graph and extract the largest connected component to define the valid cortical mantle, followed by a hole-filling step that removes spurious cortical fragments in the medial region and generates a topologically closed cortex mask. Visual inspection of the PRIME-DE dataset^5^ confirmed that this algorithm produces anatomically correct cortical masks with no obvious failures.

### Multi-contrast Pial Surface Deformation

The pial surface is generated by iteratively expanding the white matter surface toward the cerebrospinal fluid (CSF) boundary. Accurate placement is particularly challenging in deep sulcal regions where gray–CSF contrast in T1w images is often weak, and in high-field acquisitions where vascular structures can confound cortical boundaries. To improve robustness, we adopt an iterative, multi-contrast pial surface reconstruction strategy inspired by the Human Connectome Project pipelines^16^, in which different contrast cues are progressively integrated to guide surface placement.

In the initial iteration, CSF information derived from the aseg volume is used as a soft constraint to modulate local T1w intensity profiles, encouraging the evolving pial surface to expand sufficiently into sulcal depths. Subsequent refinement iterations incorporate T2w contrast (when available) or a synthetic T2w-like contrast generated from T1w via intensity inversion to fine-tune pial surface placement.

Through this multi-stage refinement process, multiple candidate pial surfaces are generated, reflecting different trade-offs between sensitivity to sulcal depth and resistance to noise and vascular artifacts, enabling flexible selection of the most anatomically plausible surface for downstream analysis.

#### Surface-aware Volumetric Registration

Volume-based registration and smoothing can degrade the spatial localization of cortical areas, as cortical fields separated along the folded sheet may lie close together in Euclidean volume space^23,55,56^. To address this, MacaSurfer implements Surface-Aware Volumetric Registration within the FireANTs framework^45^, which augments the symmetric diffeomorphic registration objective^32^ with a surface-aware loss term that penalizes the distance between deformed subject and template cortical surfaces, jointly constraining the deformation field to align cortical geometry while maintaining volumetric smoothness. Full algorithmic details and the multi-task loss formulation are provided in Supplementary Note 5.

### Normative Modelling of Macaque Brain Morphology

Using precisely reconstructed cortical and subcortical features, we built population-level normative models to enable individualized anatomical assessment. These models were derived from a multi-site cohort of 835 neurologically healthy macaques (1,145 imaging sessions, 26 sites), predominantly rhesus macaques (Macaca mulatta; number of sessions, n = 809) and a smaller subset of cynomolgus macaques (Macaca fascicularis; number of sessions, n = 26), spanning ages from birth to 23 years.

All T1-weighted scans were processed with MacaSurfer v3.0 to generate cortical surface reconstructions and subcortical segmentations. Regional morphometric measures were extracted using three cortical parcellation schemes—M129 (91 regions per hemisphere), M132 (132 regions per hemisphere)^57^, and MacBNA124 (124 regions per hemisphere)^14^—together with bilateral subcortical labels from the SARM atlas^29^. For each cortical region we quantified cortical thickness, surface area, gray matter volume, mean curvature, and sulcal depth; subcortical structures were characterized by regional volume.

Morphometric measures were harmonized across sites via the MEBRAINS framework using ComBat-GAM^27^ to reduce site-related variance while preserving biological effects, with imaging site treated as a batch variable and age, sex, and species retained during harmonization^58,59^. Regional normative trajectories were estimated using Bayesian linear regression implemented in PCNtoolkit^60^, with sex and species included as fixed-effect covariates and feature-specific global measures as nuisance regressors. Age-related trajectories were modeled with cubic B-spline basis functions, and heteroscedastic likelihoods were retained when they improved cross-validated predictive performance^61^. Full details of hyperparameter optimization and model selection are provided in Supplementary Result 1.

### Standard Output

To support transparent and efficient validation of pipeline performance, MacaSurfer produces visual inspection reports for every key processing step. These reports enable researchers to quickly identify anomalies and verify the fidelity of intermediate and final results. Visual inspection reports were generated for the following stages: raw data, skull stripping, AC-PC alignment, brain mask editing, cross-modal registration, template normalization, tissue segmentation, bias field correction, cortical reconstruction, and surface-aware volumetric registration. An example visual report is provided in Supplementary Note 6.

### Visual Quality Control (QC) Reports

MacaSurfer produces visual inspection reports for every key processing step, including skull stripping, tissue segmentation, bias field correction, cortical reconstruction, and surface-aware registration. An example report is provided in Supplementary Result 4.

## Materials

### Data collection

We aggregated structural MRI data from 39 international imaging sites, totaling 2,366 scans from 966 subjects. After processing and manual quality inspection, one subject (sub-1072, site-uwmadison) was excluded owing to empty NIfTI volumes, resulting in a final curated dataset of 2,365 scans from 965 subjects. Site-level details are provided in Supplementary Table 8.

### Normative Modelling

To construct normative models, we performed a comprehensive characterization of the dataset across age, sex, site, body weight, and breed. For subjects with longitudinal data, scans acquired at different age points were treated as independent samples to maximize the model’s representational capacity. Normative models were built from a multi-site cohort of 835 neurologically healthy macaques (from the full set of 965 subjects) with complete age, sex, and breed annotations, comprising 1,145 imaging sessions (of 1,372 total sessions) acquired across 26 international sites. Site-level details for the normative subset are provided in Supplementary Table 9.

### Evaluation Tasks

To systematically verify the stability, accuracy, and generalizability of MacaSurfer, we conducted targeted validation experiments across core functional modules, covering deep learning segmentation, robustness testing, reproducibility assessment, and performance validation of key algorithmic components. Detailed dataset compositions, selection criteria, and experimental designs for each validation experiment are provided in Supplementary Evaluation.

## Author Contributions

Y.Wei. performed the experimental studies and drafted the manuscript; Y.Wei., H.Wang., C.Chu., and L.Fan. revised the manuscript; L.Chen., Y.Wang, J.Gao., L.Cheng. C.Gao. contributed by collecting and preprocessing the neuroimaging data; T.Xu. contributed analytic tools; T.Xu., H.Wang., T.Jang., Q.Zhu., C.Chu., W.Vanduffel., and L.Fan. provided theoretical guidance and informed interpretation of the results; W.Vanduffel., H.Wang., and L.Fan. supervised the overall work.

