## Supplemental File for "MacaSurfer: unified surface-volume mapping of the macaque brain across the lifespan"

### **Supplementary Note 1: Automated orientation correction for macaque MRI**

Incorrect image orientation headers are a common issue in non-human primate (NHP) MRI, and are particularly prevalent in macaque datasets. These errors often manifest as inconsistent image orientation upon loading, left–right inversion, or anterior–posterior/superior–inferior flips, and can substantially compromise the stability and reliability of downstream registration, segmentation and atlas-mapping procedures. The high frequency of such errors in macaque MRI is largely attributable to the lack of standardized acquisition conventions. Unlike human MRI, macaque imaging often relies on custom head-fixation devices, non-standard animal coils and non-supine positioning (for example, sphinx or lateral orientations)^1–3^. As a result, image orientation may be incorrectly recorded owing to scanner-specific settings, inconsistencies in sequence configuration, or omission of manual orientation adjustment during acquisition. In addition, some acquisition systems encode image orientation relative to the animal body axis rather than the conventional neuroimaging RAS (right–anterior–superior) coordinate system, further increasing orientation ambiguity.

To address this issue, we developed an automated orientation-correction procedure that restores the correct spatial orientation of macaque MRI by alignment to a standard template. The method was implemented using FSL’s FLIRT^4,5^ and was designed to recover the gross rotational agreement between each input image and an orientation-correct reference template. Because this procedure involves cross-subject registration, skull-stripped images were used to reduce the influence of non-brain tissues on registration robustness^6^. To improve computational efficiency, each input volume was first downsampled to 0.6-mm isotropic resolution, preserving only coarse anatomical structure for rapid alignment.

Registration was performed using a two-stage search strategy. A coarse search was first carried out with a 30° angular interval to identify the approximate rotational configuration, followed by a finer search with a 10° step size for local optimization. The search space covered all combinations of the three Euler angles (pitch, roll and yaw) over the range from -180° to +180°, and the rotation yielding the highest mutual information with the reference template was selected as the optimal orientation estimate.

Template-based registration alone, however, is insufficient to guarantee correct left–right labeling. Registration algorithms assume spatial correspondence between the moving image and the reference in physical space. If the input image is stored with a left–right inversion in voxel space, geometric alignment to a correctly oriented template may still produce an anatomically aligned image with incorrect hemisphere labels. To prevent this failure mode, we introduced an additional preprocessing step before registration to harmonize the voxel-space left–right orientation of the template and the target image. Specifically, after reading the header information of the input image, the reference template was mirrored or axis-reordered as needed so that its voxel coordinate orientation matched that of the target image. Registration was then performed in a voxel space with consistent left–right ordering, thereby avoiding erroneous hemisphere exchange. After orientation correction, images were converted to the standard RAS convention to ensure a consistent coordinate system for all subsequent processing steps.

This automated orientation-correction module is applicable to multiple MRI contrasts, including T1-weighted and T2-weighted images, and was successfully deployed across macaque datasets from all PRIME-DE acquisition sites. In practice, it substantially improved the robustness and level of automation of the overall preprocessing workflow.

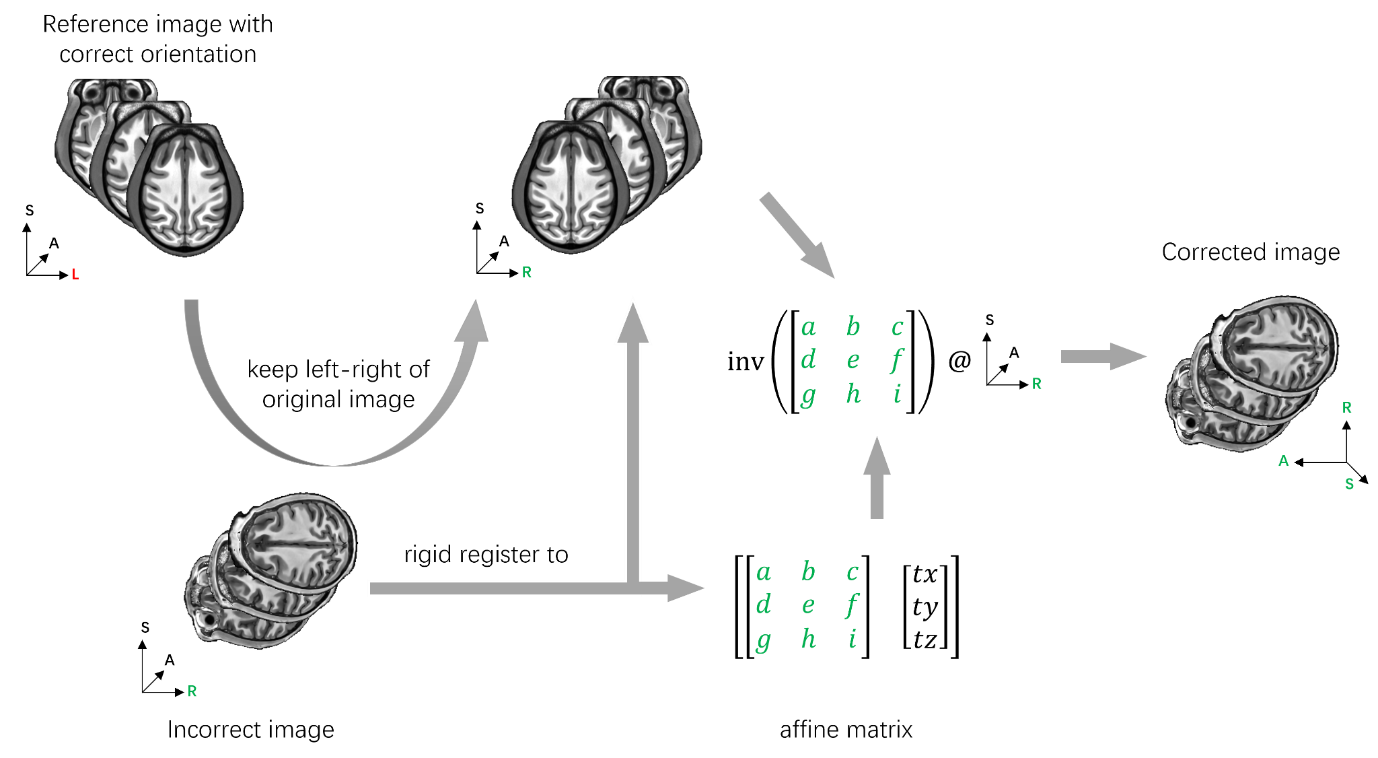

**Supplementary Figure 1.** Orientation Correction Workflow The basic idea is to use rigid registration to a standard template to obtain a rotation matrix that aligns the original image to the correct orientation. This rotation matrix is then used to extract the correct orientation. To prevent left-right hemisphere flipping, we first modify the template’s left-right orientation to match the uncorrected image before registration.

### **Supplementary Note 2: Rigid Registration Benchmarking for the MacaSurfer Pipeline**

#### **Experimental Design**

To systematically benchmark rigid registration performance for macaque structural MRI and select optimal tools for the MacaSurfer pipeline, we evaluated three widely used open-source registration packages: FSL’s FLIRT^4,5^, NiftyReg’s reg_aladin^7^, and ANTs’ antsRegistration^8^. Since rigid alignment is used for two core steps in MacaSurfer—within-subject multi-run averaging and cross-modal T1w-T2w alignment—we benchmarked all tools in both intra-modal and cross-modal registration settings:

**1. Intra-modal T1w-to-T1w registration:** 182 within-subject registration pairs from 56 randomly selected PRIME-DE subjects with multiple T1-weighted scans, registered to the first run of the first session for each subject

**2. Intra-modal T2w-to-T2w registration:** 14 within-subject registration pairs from 2 subjects with multiple T2-weighted scans

**3. Cross-modal T2w-to-T1w registration:** 232 within-subject registration pairs from 48 randomly selected subjects with paired T1w and T2w scans

All experiments were restricted to 6 degrees of freedom (3 translational, 3 rotational) consistent with rigid alignment constraints.

#### **Tool Parameter Configurations**

To minimize bias from tool-specific default settings, multiple parameter configurations were tested for each package, as detailed in Supplementary Table 1:

**Supplementary Table 1. Tested parameter configurations for rigid registration tools**

| **Registration Tool** | **Version** | **Parameter ID** | **Full Configuration Details** |
| --- | --- | --- | --- |
| FSL FLIRT | v6.0.6 | param_0 | Rigid registration (-dof 6), spline interpolation (-interp spline), default correlation ratio cost function |
| FSL FLIRT | v6.0.6 | param_1 | Rigid registration (-dof 6), spline interpolation, mutual information cost function (-cost mutualinfo) |
| FSL FLIRT | v6.0.6 | param_2 | Rigid registration (-dof 6), spline interpolation, normalized mutual information cost function (-cost normmi) |
| NiftyReg reg_aladin | v1.3.9 | param_0 | Rigid-only registration (-rigOnly), multithreading enabled (-omp 64), default zero padding (-pad 0.0) |
| NiftyReg reg_aladin | v1.3.9 | param_1 | Rigid-only registration (-rigOnly), multithreading enabled, contrast-enhanced configuration (-pad 0.0 -pv 75 -pi 75; adjusts voxel intensity percentiles for intensity mismatch robustness) |
| ANTs antsRegistration | v2.3.5 | param_0 | Rigid transform (-t Rigid[0.1]), mutual information metric (-m MI[fixed,moving,1,32]), center-of-mass initialization (-r [fixed,moving,1]), 3-level multi-resolution pyramid (--convergence [1000x500x250,1e-8,10], --shrink-factors 8x4x2, --smoothing-sigmas 4x2x1) |
| ANTs antsRegistration | v2.3.5 | param_1 | Rigid transform, mutual information metric, center-of-mass initialization, 4-level multi-resolution pyramid (--convergence [1000x500x250x100,1e-8,10], --shrink-factors 8x4x2x1, --smoothing-sigmas 3x2x1x0vox) |

#### **Evaluation Metrics**

Registration performance was evaluated using both visual inspection and quantitative metrics:

**1. For intra-modal registration:** Peak signal-to-noise ratio (PSNR) and structural similarity index (SSIM) were used as primary quantitative metrics^9^, supplemented by visual quality control^10^. All intra-modal registrations passed visual inspection.

**2. For cross-modal registration:** Direct intensity-based similarity metrics are less informative due to inherent contrast differences between T1w and T2w images, so performance was primarily evaluated via blinded visual pass rate, scored by two independent raters.

### **Supplementary Note 3: Training Details of Macaque Brain Tissue Segmentation Model**

Accurate and robust tissue segmentation is a prerequisite for cortical surface reconstruction and downstream neuroanatomical analysis. We developed a 3D Swin UNETR-based deep learning model^11,12^ for macaque brain tissue segmentation, pre-trained on large-scale non-human primate MRI data via self-supervised learning, achieving state-of-the-art performance across diverse acquisition protocols and age groups.

#### **Model Architecture**

The segmentation model is based on the 3D Swin UNETR v2 architecture, a hybrid CNN-transformer model optimized for 3D medical image segmentation. The model takes 0.4mm isotropic MRI scans as input (single channel) and outputs voxel-wise segmentation probabilities for 19 tissue classes. Key architecture parameters:

**1. Input patch size:** 96 × 96 × 96 voxels

**2. Patch size for Swin transformer:** 2 × 2 × 2

**3. Feature size:** 48

**4. Output channels:** 19 (corresponding to 18 tissue classes + background)

**Pre-training Initialization.** The model was initialized with weights from the VoCo (Volume Contrastive) self-supervised pre-trained model^13^, volume contrastive framework for 3D medical image representation learning. This pre-training provides the model with strong generalizability to anatomical variations and imaging artifacts common in MRI data. The model uses multi-scale deep supervision during training, with loss computed at 5 different resolution levels of the UNet decoder, weighted as [0.96, 0.01, 0.01, 0.01, 0.01] to prioritize the highest resolution output.

**Dataset and Segmentation Classes.** The model was trained on a large-scale multi-center macaque brain MRI dataset comprising 2,157 scans (1,699 T1-weighted, 425 T2-weighted and 33 FLAIR images) from 39 different research institutions, including both public datasets and in-house acquisitions. Four public datasets from OpenNeuro (site-ds001875^14^, site-ds003989^15^, site-ds004620^16^, site-ds005521^17^) were reserved as an independent cross-site test set for final model evaluation. The remaining data from 35 institutions were used for model training and hyperparameter selection, with an 8:2 stratified split into training and validation sets performed within each site to avoid data leakage and ensure domain consistency.

The model predicts 18 tissue and anatomical region classes, consistent with standard FreeSurfer labeling conventions for macaques:

**Supplementary Table 2. Label indices and corresponding anatomical names for macaque brain tissue segmentation**

| **Label ID** | **Region Name** |
| --- | --- |
| 0 | Background |
| 2 | Cortical gray matter |
| 3 | Cortical white matter |
| 4 | Cerebrospinal fluid (CSF) |
| 7 | Cerebellar gray matter |
| 8 | Cerebellar white matter |
| 10 | Thalamus |
| 11 | Caudate nucleus |
| 12 | Putamen |
| 13 | Globus pallidus |
| 16 | Hippocampus |
| 17 | Amygdala |
| 18 | Nucleus accumbens |
| 24 | Brainstem |
| 26 | Pons |
| 27 | Midbrain |
| 28 | Medulla oblongata |
| 138 | Substantia nigra |
| 140 | Subthalamic nucleus |

Class weights were applied during training to account for class imbalance, with higher weights assigned to small subcortical structures (gray matter, white matter, and brainstem weighted ×2 relative to larger structures).

#### **Data Preprocessing and Augmentation**

All input scans were resampled to 0.4mm isotropic resolution and intensity-normalized using percentile clipping (1st–99th percentile) prior to training. During training, the following online data augmentation strategies were applied to improve generalization:

**1. Random spatial transformations:** random axial flip, ±15° rotation, elastic deformation, and scaling (0.8–1.2×)

**2. Intensity augmentations:** random gamma correction (0.7–1.3), Gaussian and Rician noise injection, and brightness perturbation

**3. Dynamic class-balanced patch sampling:** 96×96×96 voxels patches were sampled such that under-represented small subcortical structures were sampled with higher probability, reducing training bias toward large structures.

#### **Loss Function**

Training used a combined Dice-Focal^18,19^ loss to balance segmentation accuracy across classes of different sizes:

$$\text{L}=\lambda_{\text{dice}}\text{L}_{\text{Dice}}+\lambda_{\text{focal}}\text{L}_{\text{Focal}}$$

where:

$\text{L}_{\text{Dice}}$ is the soft Dice loss, measuring overlap between predicted and ground truth segmentations

$\text{L}_{\text{Focal}}$ is the focal loss with $\gamma=3.0$ and $\alpha=0.8$, down-weighting easy examples and prioritizing hard-to-classify boundary voxels

Weighting parameters: $\lambda_{\text{dice}}=1.0$, $\lambda_{\text{focal}}=1.0$

#### **Optimization Configuration**

Optimizer: AdamW^20^ with weight decay ${10}^{-5}$

Learning rate: Initial learning rate $2\times{10}^{-3}$

Distributed training: 4 × NVIDIA A40 GPUs using DDP (Distributed Data Parallel)

Gradient accumulation: 12 steps per gradient update, effective batch size of 72 patches per iteration

Maximum epochs: 1000

### **Supplementary Note 4: Two Complementary Bias Field Correction Algorithms for Distinct Intensity Inhomogeneity Scenarios**

The spatial frequency characteristics of MRI bias fields vary substantially across acquisition protocols, field strengths, and scanner hardware. To address this heterogeneity, we developed two independent, complementary bias field correction algorithms that users can select flexibly based on the specific bias properties of their datasets: 1) an adaptive scatter RBF algorithm, departs from the B-spline smoothing strategy that underlies widely used methods such as N3 and N4ITK^21,22^. By adaptively sampling control points in regions of high intra-class residual variance and interpolating via radial basis functions with K-nearest-neighbor approximation, the method achieves substantially higher computational throughput while maintaining accuracy for smooth, low-frequency inhomogeneity. and 2) a tissue-guided GMM algorithm, extends the classical EM-based framework^23^ and subsequently refined through atlas-based tissue priors^24^ and unified generative modeling^25^. Unlike these approaches, which model the bias field using low-frequency polynomial or DCT basis functions, our method directly leverages voxel-wise tissue segmentation labels to estimate locally varying bias via Gaussian mixture modeling with signed-distance-function-based boundary handling, enabling robust correction of high-frequency artifacts that conventional methods smooth away.

#### **Adaptive Scatter RBF Bias Correction for Low-Frequency Inhomogeneity**

Suitable for conventional clinical MRI data with slowly spatially varying bias fields, and scenarios requiring fast batch processing of large cohorts. This algorithm models slow-varying low-frequency bias fields using adaptive control point sampling and accelerated radial basis function (RBF) interpolation, achieving a favorable balance between correction accuracy and computational efficiency for high-throughput processing.

Preprocessing & Adaptive Control Point Initialization

Input MRI images are first normalized to the range $[{10}^{-3},1]$ within the brain mask to ensure numerical stability during optimization. To reduce sensitivity to hard segmentation boundaries, we generate soft tissue labels by Gaussian smoothing of discrete tissue annotations:

$$w_{c}(\text{x})=\frac{G_{\sigma_{s}}(\text{I}[l(\text{x})=c])}{\sum_{c^{\text{'}}} G_{\sigma_{s}}(\text{I}[l(\text{x})=c^{\text{'}}])+\epsilon}$$

where $l(\text{x})$ is the discrete tissue label at voxel $\text{x}$, $G_{\sigma_{s}}$ is a Gaussian kernel with standard deviation $\sigma_{s}=0.5$, $\text{I}[\cdot]$ is the indicator function, and $\epsilon={10}^{-10}$ avoids division by zero.

We use a hybrid sampling strategy to initialize RBF control points: 20% of points are uniformly sampled across the brain mask for global coverage, while 80% are adaptively sampled from regions with high intra-class residual variance, which correspond to areas with stronger bias field variations. The residual variance map is computed as:

$$\mu_{c}=\frac{\sum_{\text{x}} I(\text{x})w_{c}(\text{x})}{\sum_{\text{x}} w_{c}(\text{x})+\epsilon}, V(\text{x})=\sum_{c} w_{c}(\text{x}){(I(\text{x})-\mu_{c})}^{2}$$

where $I(\text{x})$ is the normalized input image intensity, and $\mu_{c}$ is the mean intensity of tissue class $c$. Local maxima are identified via 3×3×3 maximum filtering, and control points are sampled within these high-variance regions after morphological dilation. All control points are constrained to lie within brain mask bounds.

**RBF Bias Field Modeling**

The bias field $B(\text{x})$ is modeled using Gaussian RBF interpolation. To reduce computational complexity, each voxel only considers the $K$ nearest control points (typically $K=8$–48), as contributions from distant points are negligible:

$$B(\text{x})=\frac{\sum_{k\in\text{K}(\text{x})} \phi(\left\| \text{x}-\text{p}_{k} \right\|^{2})v_{k}}{\sum_{k\in\text{K}(\text{x})} \phi(\left\| \text{x}-\text{p}_{k} \right\|^{2})+\epsilon}$$

where $\text{K}(\text{x})$ is the set of $K$ nearest control points to voxel $\text{x}$, $\text{p}_{k}\in[0,1]^{3}$ and $v_{k}$ are the normalized spatial position and bias value of control point $k$, respectively. The Gaussian RBF kernel is defined as:

$$\phi(d)=exp(-\frac{d^{2}}{2\sigma\text{rbf}^{2}})$$

with kernel width $\sigma\text{rbf}$ (0.05–0.2) controlling the smoothness of the estimated bias field. This KNN interpolation reduces computational complexity from $O(NP)$ to $O(NK)$ where $N$ is the number of voxels and $P$ is the total number of control points.

**Objective Function & Joint Optimization**

We minimize a composite objective function consisting of a data fidelity term and four regularization terms to ensure stable and accurate estimation:

$$L=L_{data}+\lambda_{1}\text{R}_{bias}+\lambda_{2}\text{R}_{spacing}+\lambda_{3}\text{R}_{\text{grad}}$$

**1. Data fidelity term**

Adaptive weighted intra-class variance of the corrected image $I(\text{x})\text{/}B(\text{x})$, with higher weights assigned to voxels with larger residuals:

$$L_{data}=\sum_{c} \sum x\omega_{c}(x)\alpha(x){(\frac{I\left( x \right)}{B\left( x \right)}-\mu_{c}^{\text{'}})}^{2}$$

where $\mu_{c}^{\text{'}}$ is the weighted mean intensity of tissue class $c$ in the corrected image, and $\alpha\left( x \right)=\frac{\sqrt{R(x)}}{\text{E}[\sqrt{R(x)}]}$ is the adaptive weight.

**2. Bias regularization**

Constrains the bias field to be close to 1 to avoid over-correction:

$$R_{bias}=\sum x{(B\left( x \right)-1)}^{2}$$

**3. Spacing regularization**

Prevents control point clustering:

$$\text{R}_{\text{spacing}}=\sum\frac{k}{\underset{k^{\text{'}}\text{≠}k}{min}\text{∣}\text{p}_{k}-\text{p}_{k^{\text{'}}}\text{∣}_{2}^{2}+\epsilon}$$

**4. Gradient regularization**

Enforces bias field smoothness:

$$\text{R}_{\text{grad}}=\text{E}[\text{∣∇}B(\text{x})\text{∣}]$$

We jointly optimize control point positions $\text{p}_{k}$ and values $v_{k}$ using the Adam optimizer (learning rates 0.001 for positions, 0.01 for values) with gradient clipping to avoid instability. The learning rate is decayed by 0.5 every 5 epochs, and optimization converges when the relative objective change is lower than ${10}^{-4}$.

This method achieves >64× speedup compared to traditional grid-based RBF methods, completing whole-brain 3D MRI correction in <5 minutes on GPU hardware with >98% correction accuracy for typical low-frequency bias fields.

#### **Tissue-Guided GMM Bias Correction for High-Frequency Inhomogeneity**

Suitable for high-field MRI data, sequences with strong gradient-induced artifacts, or datasets with significant local susceptibility-induced intensity fluctuations.

To enhance robustness against high-frequency bias fields, this tissue-guided bias field estimation method leverages tissue prior information to model locally varying inhomogeneity. The method assumes the bias field acts as multiplicative noise, and the intensity distribution of each tissue class approximates a Gaussian distribution in the corrected image, incorporating nBEST tissue segmentation results as spatial priors to preserve tissue structural information during fine-grained bias estimation.

**Model Formulation**

The input degraded image $V(\text{x})$ follows the imaging model:

$$V(\text{x})=B(\text{x})U(\text{x})+n(\text{x})$$

where $U(\text{x})$ is the corrected true image, $B(\text{x})$ is the high-frequency bias field to be estimated, and $n(\text{x})$ is additive noise, which is suppressed during preprocessing using adaptive median filtering.

We assume the intensity distribution of $K$ tissue classes in the true image $U(\text{x})$ follows a Gaussian mixture model. Instead of estimating tissue prior distribution $p(y_{j}\text{∣}\text{x})$ using empirical constants or whole-image statistics, we use soft segmentation results provided by nBEST to preserve the spatial structure and relative positional relationships of tissues. To address potential misclassifications at low-contrast boundaries in nBEST segmentation results, we apply a Gaussian kernel label smoothing strategy to the segmentation results within local voxel neighborhoods, improving tissue consistency at boundaries. This local smoothing process is equivalent to introducing a Markov random field regularization term that constrains the spatial continuity of tissue distributions.

**Bias Estimation and Optimization**

For any voxel $\text{x}_{i}$, the conditional probability density belonging to tissue class $j$ is:

$$p(y_{j}\text{∣}\text{x}_{i},\mu_{j},\sigma_{j})=\frac{1}{\sqrt{2\pi}\sigma_{j}}exp(-\frac{(U(\text{x}_{i})-\mu_{j})^{2}}{2\sigma_{j}^{2}})$$

where $\mu_{j}$ and $\sigma_{j}$ are the intensity mean and standard deviation of tissue class $j$, respectively. Combining the soft segmentation prior $p(y_{j}\text{∣}\text{x}i)$ provided by nBEST, the membership weight of voxel $\text{x}i$ to class $j$ is:

$$Wij=\frac{p(y_{j}\text{∣}\text{x}_{i},\mu_{j},\sigma_{j})p(y_{j}\text{∣}\text{x}_{i})}{\sum_{j^{\text{'}}} p(y_{j^{\text{'}}}\text{∣}\text{x}_{i},\mu_{j^{\text{'}}},\sigma_{j^{\text{'}}})p(y_{j^{\text{'}}}\text{∣}\text{x}_{i})}$$

We compute voxel-wise residuals based on membership weights, and estimate the bias field from these residuals:

$$R_{i}=\sum_{j\in\text{L}} \frac{W_{ij}(V(\text{x}_{i})-\mu_{j})}{\sigma_{j}^{2}}$$

where $\text{L}$ is the set of tissue classes. To ensure spatial smoothness of the bias field, we smooth the residuals using a Gaussian filter $F$. To address artifacts at the brain boundary caused by conventional padding/reflecting strategies, we introduce a signed distance function (SDF) $\phi$ to compute the distance from extra-cerebral voxels to the brain tissue boundary, and fill extra-cerebral regions with bias values from symmetric points across the boundary, effectively mitigating boundary effects:

$$b_{i}=\frac{[FR]_{i}}{[F\phi^{-1}\text{1}]_{i}}$$

where $\text{1}$ is an all-one vector and $\phi^{-1}$ is the inverse weight of the signed distance function.

The entire process is implemented iteratively: in each iteration, we fix the bias field and update GMM parameters $\mu_{j},\sigma_{j}$, then fix the GMM parameters and update the bias field until the change in objective function is below the convergence threshold.

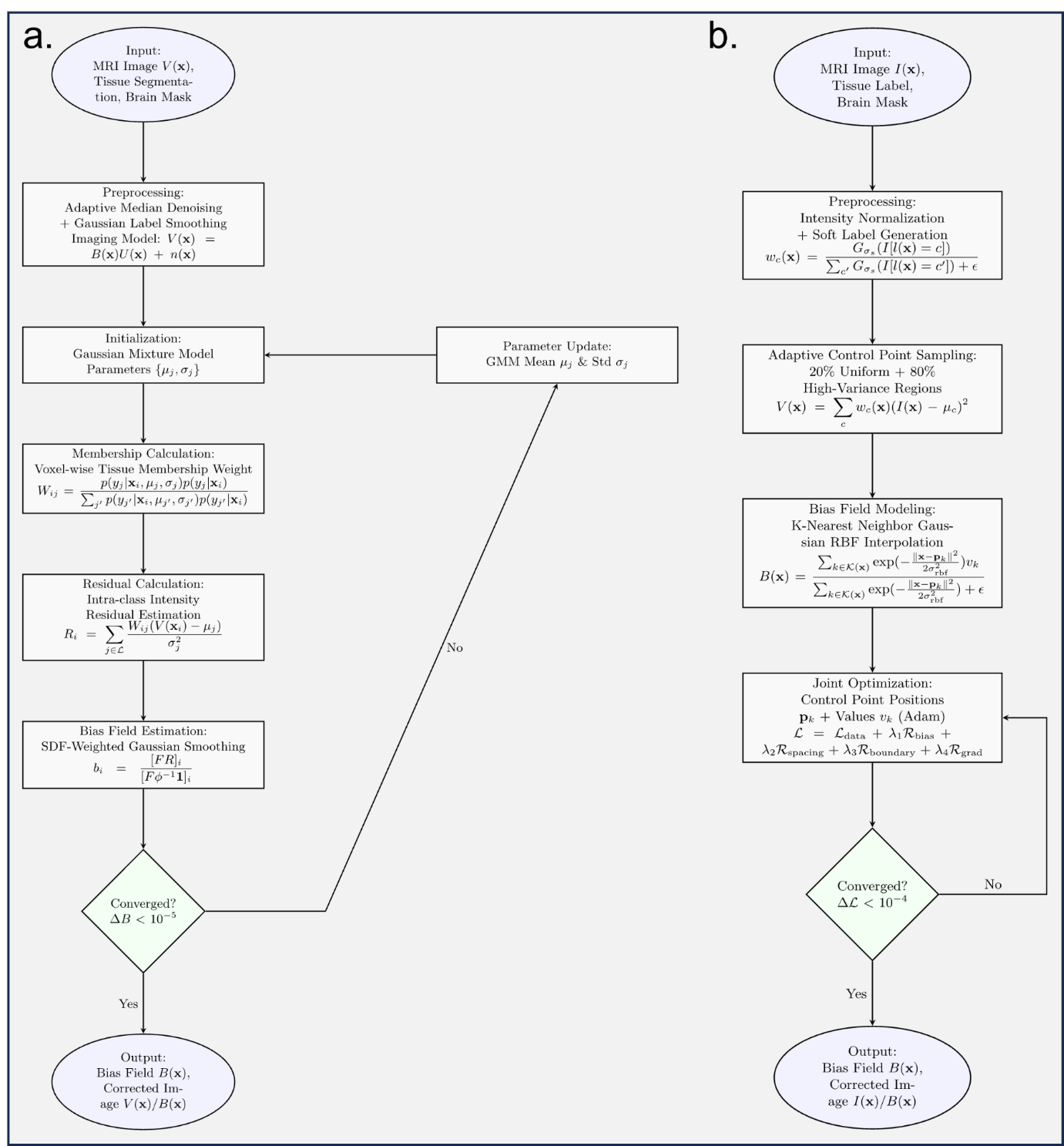

**Supplementary Figure 3.** Flowcharts of the two complementary bias field correction algorithms (a) Pipeline of the tissue-guided Gaussian mixture model (GMM) correction algorithm, optimized for robust correction of high-frequency local artifacts and susceptibility-induced intensity fluctuations; (b) Pipeline of the adaptive scatter radial basis function (RBF) correction algorithm, designed for high-throughput correction of smooth low-frequency intensity inhomogeneity.

#### **Algorithm Selection Guidance**

For most conventional T1/T2-weighted MRI datasets with slowly varying bias fields, the adaptive scatter RBF method provides optimal balance between correction accuracy and computational speed. For high-field MRI data, sequences with significant gradient artifacts, or datasets showing obvious local intensity fluctuations, the tissue-guided GMM method is preferred for its superior robustness to high-frequency inhomogeneity.

### **Supplementary Note 5: Surface-Awared Volumetric Registration for MacaSurfer**

Accurate volumetric registration is a core step in neuroimaging pipelines, but traditional intensity-only methods systematically degrade cortical spatial localization relative to surface-based approaches^26^. In human neuroimaging, the high inter-individual variability of cortical folding patterns^27^ creates a fundamental tension: aligning cortical geometry and subcortical anatomy within a single Euclidean deformation field are incompatible objectives, because the large non-local displacements required for cortical matching disrupt subcortical correspondence. Prior methods have addressed this by escaping Euclidean space—first registering cortical surfaces on a sphere, then propagating the result into the volume via harmonic mapping or elastic relaxation, followed by intensity-based refinement^28,29^. Even the most recent learning-based approaches retain a spherical intermediate space to decouple these competing constraints^30^. Macaque cortical folding, however, is far more stereotyped and shows markedly lower inter-individual variability than human folding^27^. This species-specific property fundamentally changes the registration problem: because cortical folding patterns are highly consistent across macaque individuals, surface alignment and volumetric alignment are no longer contradictory objectives in Euclidean space. We exploited this insight to develop a surface-constrained registration framework that augments the symmetric diffeomorphic objective in FireANTs^31^ with an explicit surface geometry loss term, jointly optimizing volumetric intensity similarity and cortical surface correspondence within a single Euclidean deformation field. This approach bypasses the sequential surface-then-volume pipeline necessitated by human cortical variability and instead achieves simultaneous surface-volume alignment in a single optimization step.

#### **Algorithm Framework**

The surface-constrained registration is implemented as a symmetric bidirectional deformable registration pipeline built on the FireANTs GPU-accelerated registration library^31^, with support for multi-resolution optimization and affine pre-alignment. The pipeline takes as input: (1) source and target 3D volumetric images, (2) optional corresponding cortical surface meshes (white/pial surfaces reconstructed from the same scans), and (3) optional cortical region of interest (ROI) masks for weighted alignment. The pipeline outputs the warped volume, deformation fields, and warped surface meshes if provided.

A three-level multi-resolution optimization strategy is used by default, with downsampling factors of 4×, 2×, and 1× from coarse to fine, with 600, 500, and 400 iterations per scale respectively. An initial affine alignment step is performed prior to deformable registration to correct for gross translational and rotational differences between scans.

#### **Multi-Task Loss Function**

The optimization objective combines four complementary loss terms to balance volumetric alignment accuracy, surface alignment fidelity, deformation field smoothness, and topological consistency:

$$L_{\text{total}}=\lambda_{\text{vol}}L_{\text{vol}}+\lambda_{\text{surf}}L_{\text{surf}}+\lambda_{\text{reg}}L_{\text{reg}}+\lambda_{\text{consist}}L_{\text{consist}}$$

with default weighting parameters: $\lambda_{\text{vol}}=5\times{10}^{-5}$, $\lambda_{\text{surf}}=1.0$, $\lambda_{\text{reg}}=5\times{10}^{-2}$, $\lambda_{\text{consist}}=1.0$.

**1. Volumetric Similarity Loss**

Volumetric alignment is measured using local normalized cross-correlation (NCC) with a 5×5×5 kernel, which is robust to intensity inhomogeneity and contrast differences between scans:

$$\text{L}_{\text{vol}}=\text{L}_{\text{NCC}}(I_{\text{moving}}\circ\phi_{\text{fwd}},I_{\text{fixed}})+\text{L}_{\text{NCC}}(I_{\text{fixed}}\circ\phi_{\text{rev}},I_{\text{moving}})$$

where $\phi_{\text{fwd}}$ and $\phi_{\text{rev}}$ are the forward and reverse deformation fields, respectively, and $\circ$ denotes warping via spatial transformation. The loss is computed symmetrically for both forward (moving→fixed) and reverse (fixed→moving) directions to ensure topological consistency.

**2. Surface Alignment Loss**

When cortical surface meshes are provided, we add a geometric alignment loss that penalizes distance between corresponding vertices of the warped source surface and target surface. The loss supports three distance metrics: mean squared error (MSE), L1 error, and Chamfer distance, with MSE used as the default for vertex-corresponding surfaces:

$$\text{L}_{\text{surf}}=\text{L}_{\text{mesh}}(\phi_{\text{fwd}}(S_{\text{fixed}}),S_{\text{moving}})+\text{L}_{\text{mesh}}(\phi_{\text{rev}}(S_{\text{moving}}),S_{\text{fixed}})$$

where $S_{\text{fixed}}$ and $S_{\text{moving}}$ are the fixed and moving surface vertex coordinates, respectively, and $\phi_{\text{fwd}}(S)$ denotes warping of surface vertices via the deformation field. For MSE loss:

$$\text{L}_{\text{mesh}}(S_{\text{warped}},S_{\text{target}})=\frac{1}{N}\sum_{i=1}^{N} w_{i}{\parallel\frac{S_{\text{warped},i}}{\text{V}}-\frac{S_{\text{target},i}}{\text{V}}\parallel}_{2}^{2}$$

where $\text{V}$ is the volume dimension normalization factor, and $w_{i}$ is the ROI weight (set to 1 for cortical vertices, 0.2 for non-cortical vertices by default to prioritize alignment of gray matter regions). For non-corresponding surfaces, Chamfer distance is used to measure the average nearest-neighbor distance between two-point clouds.

**3. Deformation Regularization Loss**

A diffusion regularization term is applied to the displacement field to enforce spatial smoothness and prevent unrealistic local deformations:

$$\text{L}_{\text{reg}}=\text{∥∇}\phi_{\text{fwd}}\text{∥}_{2}^{2}+\text{∥∇}\phi_{\text{rev}}\text{∥}_{2}^{2}$$

where $\text{∇}\phi$ denotes the spatial gradient of the displacement field.

**4. Cycle Consistency Loss**

A bidirectional cycle consistency loss ensures that the forward and reverse deformation fields are approximately inverses of each other, improving topological correctness:

$$\text{L}_{\text{consist}}=\text{∥}\phi_{\text{fwd}}\circ\phi_{\text{rev}}-\text{Id}\text{∥}_{2}^{2}+\text{∥}\phi_{\text{rev}}\circ\phi_{\text{fwd}}-\text{Id}\text{∥}_{2}^{2}$$

where $\text{Id}$ is the identity transformation.

#### **Optimization Strategy**

All parameters are optimized using the AdamW^20^ optimizer with a learning rate of 0.4, decayed automatically across multi-resolution scales. The optimization converges when the relative change in total loss is less than ${10}^{-12}$ for 20 consecutive iterations. The entire pipeline is GPU-accelerated, completing a typical 3D macaque MRI registration in <5 minutes on an NVIDIA A40 GPU.

### **Supplementary Note 6: MacaSurfer Standard Output Directory Structure**

Following the completion of cortical reconstruction, normative modeling, and all upstream processing workflows, MacaSurfer generates structured, standardized outputs organized by functional stage to ensure traceability and support reproducible downstream analyses. Preprocessed structural MRI data are systematically organized into five primary directories, each corresponding to a specific processing stage:

**Prepare Directory:** This directory stores run-level preprocessing outputs under user-defined or default configurations, including skull-stripped volumes, corrected anatomical coordinates, refined brain masks, rigid-body alignment across multiple runs, and averaged denoised volumes.

**Enhance Directory:** This directory contains results from advanced volumetric processing, including registration to standard templates, tissue segmentation, refined white matter masks, ACPC alignment, and tissue-guided bias field correction.

**Surface Directory:** This directory holds the white and pial cortical surfaces reconstructed by FreeSurfer, together with intermediate files generated during the surface reconstruction procedure.

**Resample Directory:** This directory provides surfaces resampled to multiple coordinate spaces (including native image space, AC-PC aligned space, and nonlinearly transformed template space). For each space, surfaces are available at three vertex resolutions (original vertex count, 32k, and 164k), accompanied by parcellation files, cortical thickness maps, curvature maps, and sulcal depth maps.

**QC Directory:** This directory contains visual quality-control reports for each processing stage as well as statistical summaries derived from normative modeling.

This directory structure ensures that both intermediate and final outputs are clearly organized and readily accessible for quality assessment and downstream analyses.

### **Supplementary Result 1: Details of Normative modelling for non-human primate brain morphometry**

#### **Normative model performance**

**Supplementary Table 3. Normative model performance across morphometric modalities**

| **Modality** | **Mean EXPV ± s.d.** | **Mean MAE ± s.d.** | **Mean RMSE ± s.d.** | **Mean R² ± s.d.** | **Mean Spearman's ρ ± s.d.** |
| --- | --- | --- | --- | --- | --- |
| Cortical thickness | 0.24 ± 0.10 | 0.020 ± 0.009 mm | 0.17 ± 0.06 mm | 0.24 ± 0.10 | 0.39 ± 0.10 |
| Gray matter area | 0.22 ± 0.07 | 0.018 ± 0.009 mm² | 16.3 ± 30.0 mm² | 0.22 ± 0.07 | 0.40 ± 0.08 |
| Gray matter volume | 0.25 ± 0.09 | 0.018 ± 0.010 mm³ | 35.8 ± 59.2 mm³ | 0.25 ± 0.09 | 0.41 ± 0.08 |
| Cortical curvature | 0.06 ± 0.06 | 0.038 ± 0.028 | 0.03 ± 0.05 | 0.06 ± 0.06 | 0.22 ± 0.10 |
| Sulcal depth | 0.11 ± 0.06 | 0.024 ± 0.013 | 0.81 ± 0.26 | 0.11 ± 0.06 | 0.29 ± 0.10 |
| Subcortical volume | 0.39 ± 0.12 | 0.017 ± 0.004 mm³ | 284.7 ± 606.8 mm³ | 0.39 ± 0.12 | 0.58 ± 0.11 |

*MAE = mean absolute error, RMSE = root mean squared error, R² = coefficient of determination, ρ = Spearman correlation coefficient. All metrics are reported as cross-validated out-of-sample estimates on the held-out test set.*

*Abbreviations: EXPV, explained variance; s.d., standard deviation. Performance metrics are reported as cross-validated out-of-sample estimates.*

#### **Regional and parcellation effects**

**Supplementary Table 4. Regional variation in model performance (cortical thickness, SA124 atlas, left hemisphere)**

| **Brain region** | **EXPV** | **Spearman's ρ** | **P value** |
| --- | --- | --- | --- |
| Inferior temporal gyrus (ITG) | 0.52 | 0.57 | < 1 × 10⁻⁹⁹ |
| Cingulate gyrus (ventral rostral) | 0.45 | 0.56 | < 1 × 10⁻⁹³ |
| Orbital gyrus (rostromedial) | 0.43 | 0.57 | < 1 × 10⁻¹⁰¹ |
| Precuenus (dorsal intermediate) | 0.42 | 0.58 | < 1 × 10⁻¹⁰¹ |
| Ventromedial prefrontal cortex | 0.42 | 0.59 | < 1 × 10⁻¹⁰⁶ |
| Temporal pole (ventral rostral) | 0.02 | 0.16 | < 1 × 10⁻⁷ |
| Insular cortex (ventral rostral) | 0.04 | 0.11 | < 1 × 10⁻³ |
| Temporal pole (lateral rostral) | 0.04 | 0.20 | < 1 × 10⁻¹¹ |

Performance was consistent across the three cortical parcellation schemes, with average EXPV ranging from 0.25 ± 0.19 (MBNA124 atlas) to 0.28 ± 0.22 (M129 atlas). The multi-modal combined atlas (Modalities) achieved the highest average performance (0.41 ± 0.32), driven by its superior surface area and thickness model performance.

#### **Hyperparameter optimization results**

All model hyperparameters were rigorously optimized via 5-fold cross-validation over a grid of 32 parameter combinations, ensuring that each model was tuned for optimal out-of-sample generalization. The final model ensemble used the top-performing parameters for each modality and brain region, with all models trained on the full dataset for downstream applications. The complete optimal hyperparameter selections and cross-validation performance for all models are provided in Supplementary Table 4 below.

**Supplementary Table 5. Optimal hyperparameters and cross-validation performance for all normative models**

| **Atlas** | **Hemisphere** | **Modality** | **Number of knots** | **Spline degree** | **Heteroskedastic noise** | **Adaptive knot placement** | **Mean EXPV (cross-validation)** | **Std EXPV (cross-validation)** |
| --- | --- | --- | --- | --- | --- | --- | --- | --- |
| MBNA124 | L | Cortical curvature | 2 | 2 | True | True | 0.0656 | 0.0223 |
| MBNA124 | L | Gray matter area | 2 | 3 | False | True | 0.4612 | 0.0109 |
| MBNA124 | L | Gray matter volume | 2 | 3 | False | True | 0.4274 | 0.0157 |
| MBNA124 | L | Cortical thickness | 2 | 3 | True | True | 0.2716 | 0.0145 |
| MBNA124 | L | Sulcal depth | 2 | 3 | False | True | 0.0685 | 0.0032 |
| MBNA124 | L | Global mean thickness | 2 | 2 | False | True | 0.0589 | 0.0477 |
| MBNA124 | L | Global mean sulcal depth | 2 | 2 | False | True | 0.2714 | 0.0384 |
| MBNA124 | L | Global mean curvature | 2 | 2 | True | False | 0.2138 | 0.0972 |
| MBNA124 | L | Global total gray matter area | 2 | 3 | False | True | 0.1963 | 0.0344 |
| MBNA124 | L | Global total cortical volume | 4 | 2 | True | False | 0.1220 | 0.0282 |
| MBNA124 | R | Sulcal depth | 4 | 3 | True | False | 0.0579 | 0.0090 |
| MBNA124 | R | Gray matter area | 2 | 3 | False | True | 0.4637 | 0.0140 |
| MBNA124 | R | Cortical thickness | 2 | 3 | False | True | 0.2836 | 0.0234 |
| MBNA124 | R | Cortical curvature | 2 | 2 | True | False | -0.0124 | 0.0822 |
| MBNA124 | R | Gray matter volume | 2 | 3 | False | True | 0.4323 | 0.0093 |
| MBNA124 | R | Global mean curvature | 2 | 3 | True | False | 0.0526 | 0.0585 |
| MBNA124 | R | Global mean sulcal depth | 2 | 2 | True | True | 0.2783 | 0.0331 |
| MBNA124 | R | Global mean thickness | 2 | 3 | False | True | 0.1883 | 0.0614 |
| MBNA124 | R | Global total gray matter area | 2 | 3 | False | True | 0.2046 | 0.0315 |
| MBNA124 | R | Global total cortical volume | 4 | 2 | True | False | 0.1219 | 0.0344 |
| M129 | L | Cortical curvature | 2 | 2 | True | True | 0.0556 | 0.0542 |
| M129 | L | Gray matter area | 2 | 3 | False | True | 0.4985 | 0.0172 |
| M129 | L | Sulcal depth | 2 | 3 | True | True | 0.0772 | 0.0077 |
| M129 | L | Gray matter volume | 2 | 3 | False | True | 0.4736 | 0.0154 |
| M129 | L | Cortical thickness | 2 | 3 | False | True | 0.3200 | 0.0198 |
| M129 | L | Global total gray matter area | 4 | 3 | False | False | 0.1601 | 0.0293 |
| M129 | L | Global mean thickness | 2 | 3 | True | True | 0.0865 | 0.0620 |
| M129 | L | Global mean curvature | 2 | 2 | False | False | 0.1611 | 0.0775 |
| M129 | L | Global mean sulcal depth | 2 | 2 | False | True | 0.2364 | 0.0463 |
| M129 | L | Global total cortical volume | 4 | 2 | False | False | 0.2169 | 0.0399 |
| M129 | R | Gray matter area | 2 | 3 | False | True | 0.5133 | 0.0078 |
| M129 | R | Gray matter volume | 2 | 3 | True | True | 0.4819 | 0.0109 |
| M129 | R | Sulcal depth | 2 | 3 | True | True | 0.0720 | 0.0052 |
| M129 | R | Cortical curvature | 2 | 3 | True | True | -0.0478 | 0.1635 |
| M129 | R | Cortical thickness | 2 | 3 | False | True | 0.3322 | 0.0274 |
| M129 | R | Global total gray matter area | 2 | 3 | True | True | 0.1194 | 0.0462 |
| M129 | R | Global mean thickness | 2 | 3 | False | True | 0.1891 | 0.0623 |
| M129 | R | Global mean curvature | 2 | 3 | False | True | 0.0052 | 0.0221 |
| M129 | R | Global mean sulcal depth | 2 | 2 | True | True | 0.2641 | 0.0691 |
| M129 | R | Global total cortical volume | 2 | 2 | False | True | 0.1911 | 0.0408 |
| M132 | R | Gray matter area | 2 | 3 | False | True | 0.4894 | 0.0172 |
| M132 | R | Cortical thickness | 2 | 3 | True | True | 0.2966 | 0.0283 |
| M132 | R | Sulcal depth | 4 | 3 | True | False | 0.0609 | 0.0041 |
| M132 | R | Cortical curvature | 2 | 2 | True | False | -0.0218 | 0.1056 |
| M132 | R | Gray matter volume | 2 | 3 | True | True | 0.4425 | 0.0101 |
| M132 | R | Global mean curvature | 3 | 2 | False | False | 0.0018 | 0.0115 |
| M132 | R | Global mean thickness | 2 | 3 | True | True | 0.1058 | 0.0748 |
| M132 | R | Global mean sulcal depth | 2 | 2 | False | True | 0.2779 | 0.0674 |
| M132 | R | Global total gray matter area | 2 | 3 | True | True | 0.1338 | 0.0262 |
| M132 | R | Global total cortical volume | 4 | 2 | True | False | 0.1195 | 0.0343 |
| M132 | L | Cortical thickness | 2 | 3 | False | True | 0.2876 | 0.0170 |
| M132 | L | Gray matter area | 2 | 3 | False | True | 0.4822 | 0.0178 |
| M132 | L | Cortical curvature | 2 | 3 | True | True | 0.0734 | 0.0272 |
| M132 | L | Gray matter volume | 2 | 3 | False | True | 0.4455 | 0.0186 |
| M132 | L | Sulcal depth | 2 | 3 | True | True | 0.0652 | 0.0076 |
| M132 | L | Global mean thickness | 4 | 2 | False | False | 0.0421 | 0.0287 |
| M132 | L | Global mean curvature | 4 | 2 | False | False | 0.1719 | 0.0990 |
| M132 | L | Global mean sulcal depth | 4 | 2 | True | False | 0.2816 | 0.0462 |
| M132 | L | Global total gray matter area | 2 | 3 | True | True | 0.1914 | 0.0495 |
| M132 | L | Global total cortical volume | 2 | 3 | False | True | 0.1165 | 0.0405 |
| Modalities | L | Gray matter area | 2 | 3 | True | True | 0.6881 | 0.0164 |
| Modalities | L | Cortical thickness | 2 | 3 | False | True | 0.5560 | 0.0205 |
| Modalities | L | Cortical curvature | 2 | 2 | False | True | 0.1091 | 0.1472 |
| Modalities | L | Sulcal depth | 2 | 3 | False | True | 0.1786 | 0.0144 |
| Modalities | L | Gray matter volume | 2 | 3 | True | True | 0.7038 | 0.0112 |
| Modalities | L | Global mean thickness | 2 | 3 | False | True | 0.0314 | 0.0336 |
| Modalities | L | Global mean curvature | 2 | 3 | False | True | 0.1550 | 0.0739 |
| Modalities | L | Global mean sulcal depth | 2 | 2 | False | True | 0.2037 | 0.0455 |
| Modalities | L | Global total gray matter area | 4 | 3 | True | False | 0.1244 | 0.0242 |
| Modalities | L | Global total cortical volume | 2 | 2 | False | True | 0.2240 | 0.0375 |
| Modalities | R | Cortical curvature | 2 | 2 | True | False | -0.2024 | 0.4978 |
| Modalities | R | Sulcal depth | 2 | 3 | False | True | 0.1589 | 0.0256 |
| Modalities | R | Gray matter area | 2 | 3 | False | True | 0.6860 | 0.0284 |
| Modalities | R | Cortical thickness | 2 | 3 | False | True | 0.5603 | 0.0315 |
| Modalities | R | Gray matter volume | 2 | 3 | True | True | 0.6831 | 0.0172 |
| Modalities | R | Global mean curvature | 2 | 3 | True | True | -0.0095 | 0.0143 |
| Modalities | R | Global mean thickness | 2 | 3 | False | True | 0.1738 | 0.0596 |
| Modalities | R | Global mean sulcal depth | 4 | 2 | False | False | 0.2772 | 0.0291 |
| Modalities | R | Global total gray matter area | 2 | 3 | True | True | 0.1283 | 0.0263 |
| Modalities | R | Global total cortical volume | 2 | 2 | False | True | 0.1873 | 0.0387 |
| Subcortical | L | Subcortical volume | 2 | 3 | False | True | 0.3909 | 0.0540 |
| Subcortical | L | Global estimated ICV | 2 | 3 | False | True | 0.3383 | 0.0765 |
| Subcortical | L | Global total subcortical volume | 2 | 3 | False | True | 0.4024 | 0.0674 |
| Subcortical | R | Subcortical volume | 2 | 3 | False | True | 0.3812 | 0.0343 |
| Subcortical | R | Global estimated ICV | 2 | 3 | False | True | 0.3203 | 0.0536 |
| Subcortical | R | Global total subcortical volume | 2 | 3 | False | True | 0.3932 | 0.0418 |

*EXPV = explained variance. All models were evaluated over 32 hyperparameter combinations with 5-fold cross-validation.*

Cross-validation identified consistent optimal hyperparameter patterns across modalities:

- 95% of models selected 2 internal B-spline knots, balancing model flexibility and generalization
- 86% of models used cubic B-splines (degree=3), with quadratic splines (degree=2) preferred for low-signal modalities including curvature and sulcal depth
- 55% of models favored homoscedastic noise structure, with heteroscedastic noise providing small improvements for surface area and thickness models
- 88% of models selected adaptive quantile-based knot placement rather than uniform spacing, better accommodating the non-uniform age distribution of the cohort

Across all models, the 5-fold cross-validation procedure achieved highly consistent performance estimates, with an average standard deviation of EXPV across folds of 0.049, indicating robust model generalizability.

#### **Covariate contribution**

Including global brain measures as covariates significantly improved model performance across all modalities (Supplementary Table 6). Compared to baseline models using only age, sex, and breed as predictors, adding global covariates increased mean EXPV by 0.22 for surface area models, 0.18 for thickness models, and 0.19 for volume models (all P < 1 × 10⁻¹⁵, paired t-test across regions).

**Supplementary Table 6. Contribution of global covariates to model performance**

| **Modality** | **Mean EXPV (baseline: age+sex+breed)** | **Mean EXPV (full model: + global covariates)** | **ΔEXPV** | **P value (paired t-test)** |
| --- | --- | --- | --- | --- |
| Cortical thickness | 0.10 ± 0.06 | 0.24 ± 0.10 | 0.14 | < 1 × 10⁻¹⁵ |
| Gray matter area | 0.08 ± 0.04 | 0.22 ± 0.07 | 0.14 | < 1 × 10⁻¹⁸ |
| Gray matter volume | 0.11 ± 0.05 | 0.25 ± 0.09 | 0.14 | < 1 × 10⁻¹⁶ |
| Cortical curvature | 0.02 ± 0.03 | 0.06 ± 0.06 | 0.04 | < 1 × 10⁻⁶ |
| Sulcal depth | 0.04 ± 0.03 | 0.11 ± 0.06 | 0.07 | < 1 × 10⁻¹⁰ |

*Values are derived from 5-fold cross-validation comparisons between models with and without global brain size covariates. All comparisons are statistically significant after false discovery rate correction.*

#### **Biological validity**

All models captured biologically plausible age-related trajectories consistent with prior primate neurodevelopment studies, with the unique advantage of our sample covering the full developmental spectrum from birth to early adulthood. Cortical thickness showed significant thinning during adolescence (slope = -0.008 mm/year, P < 0.001), stabilizing in early adulthood (~10 years of age). Gray matter area increased rapidly during postnatal development, peaking at 3–4 years of age before declining slowly through adolescence and adulthood. Curvature and sulcal depth showed weaker age effects, consistent with greater developmental stability of these morphological features established early in life. All models satisfied the normality assumption of residuals (Shapiro–Wilk test, P > 0.05 after false discovery rate correction across all regions).

The performance of our models is comparable to state-of-the-art human normative models (which typically achieve EXPV 0.4–0.6 for cortical thickness) and represents the first large-scale normative model for non-human primates, outperforming previous smaller-scale NHP models (mean EXPV = 0.31 in the only prior published study of macaque normative modeling).

#### **Data and code availability**

All trained normative models, code for preprocessing, harmonization, and model fitting, and example usage tutorials are publicly available at [https://github.com/your-lab/monkey-normative-models]

### **Supplementary Result 2: Results of Key Components in MacaSurfer**

#### **Automated Orientation Correction**

To evaluate the robustness of the automated orientation correction module, we conducted experiments across 39 imaging sites. For each site, one subject’s T1-weighted image (after skull stripping) was randomly selected and artificially assigned an incorrect orientation through random (non-mirrored) axis permutations.

The proposed orientation correction algorithm was then applied to these images. Visual inspection confirmed that all cases were successfully restored to the correct anatomical orientation across all sites.

To further quantify the impact of orientation errors, we performed affine registration (12 degrees of freedom) to the template space using NiftyReg (reg_aladin) on both the misoriented and corrected images. Registration failed in most cases when incorrect orientation was present, with only a small number of successful alignments. In contrast, orientation-corrected images consistently achieved accurate alignment.

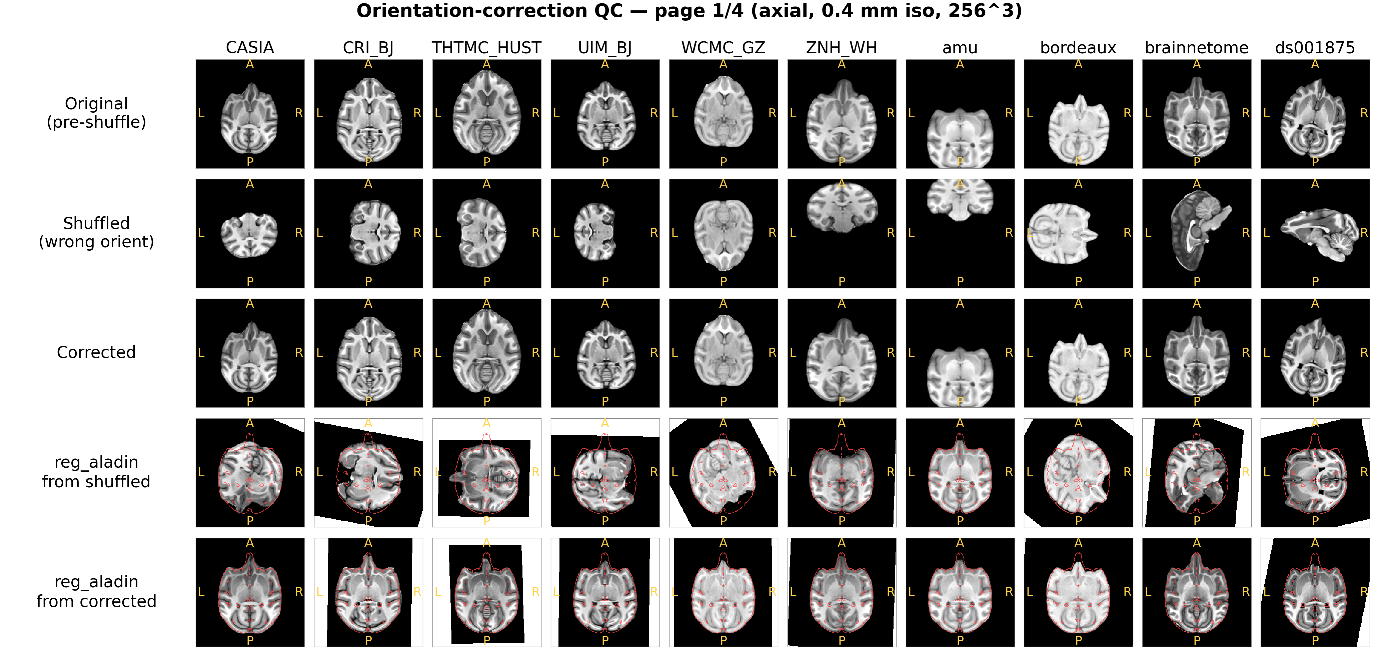

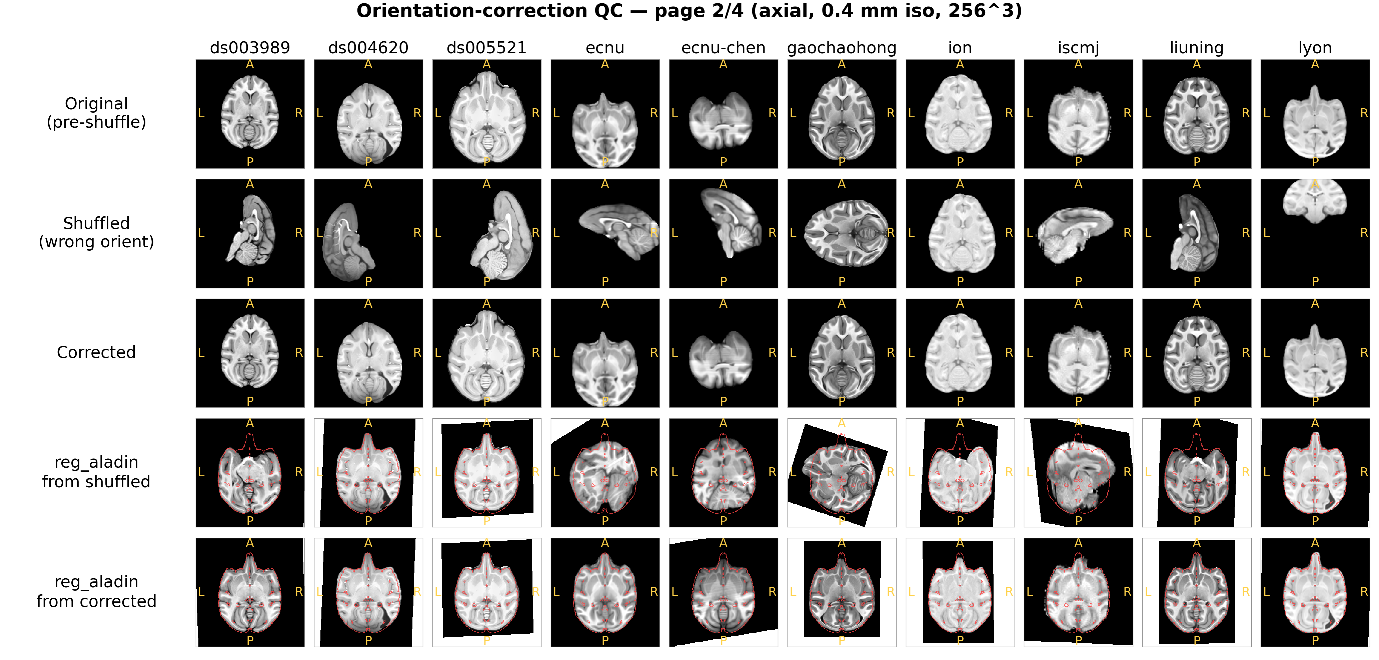

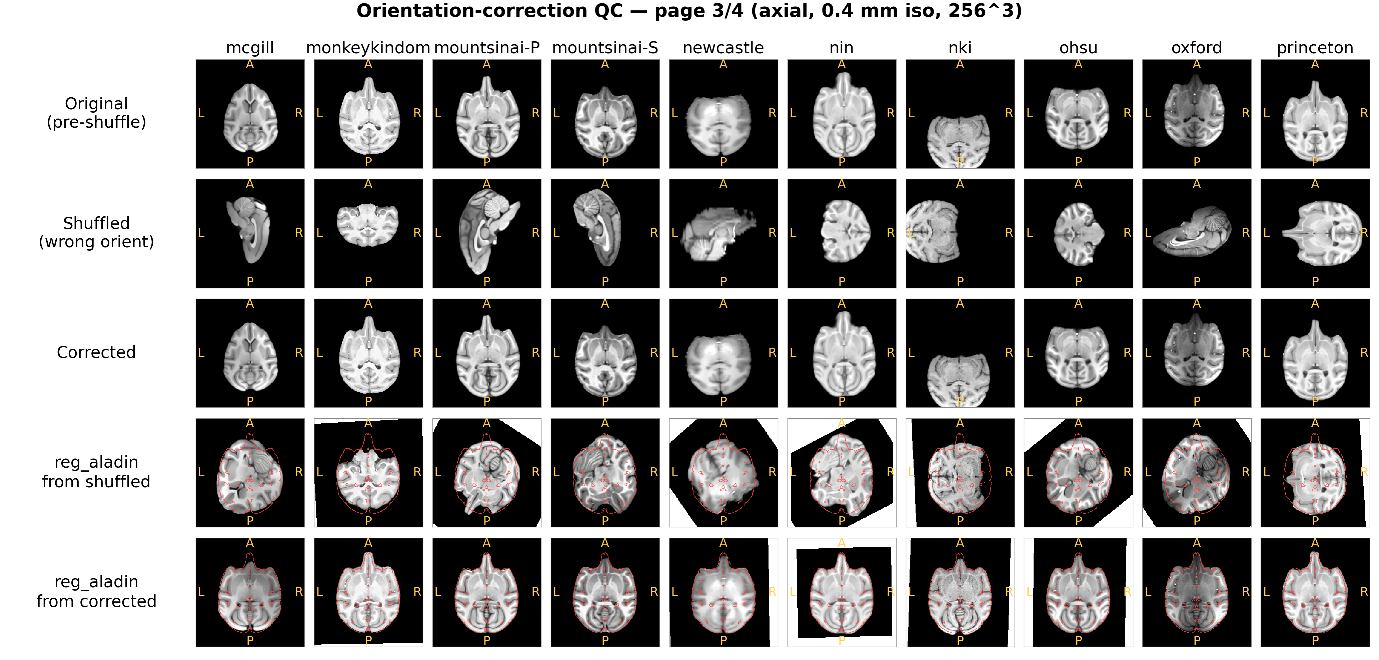

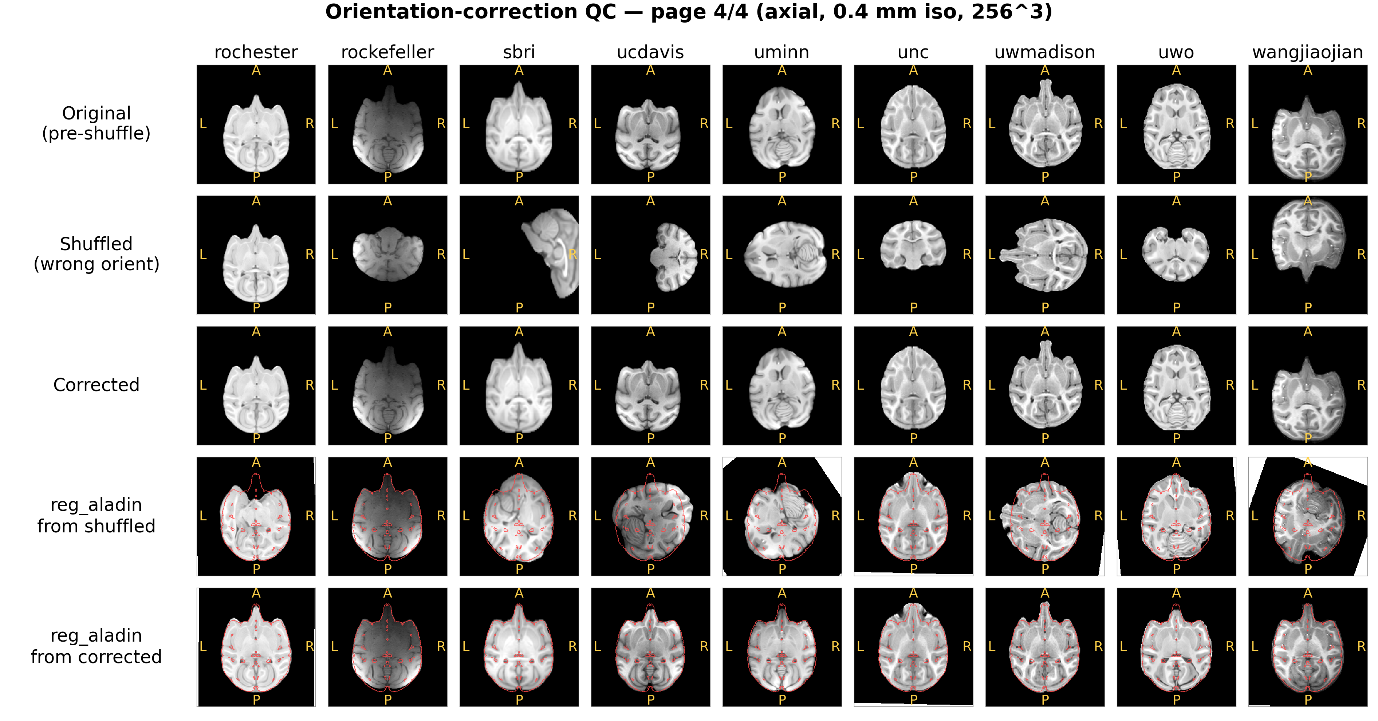

**Supplementary Figure 4. Visualization of automated orientation correction across sites.** For each of the 39 imaging sites, one subject was randomly selected. Shown are the original image, the image after random orientation perturbation, and the result after automated orientation correction. Due to the large number of sites, results are displayed across four panels. In each panel, the last two rows show the corresponding registration results of misoriented and corrected images to the MEBRAINS template. For visual assessment of alignment quality, template contours are overlaid on the images.

Quantitative evaluation was performed using normalized cross-correlation (NCC), computed within the brain mask after registration. Across nearly all sites, orientation correction led to a substantial increase in NCC values, demonstrating that incorrect anatomical orientation is a major source of registration failure and that the proposed method effectively resolves this issue.

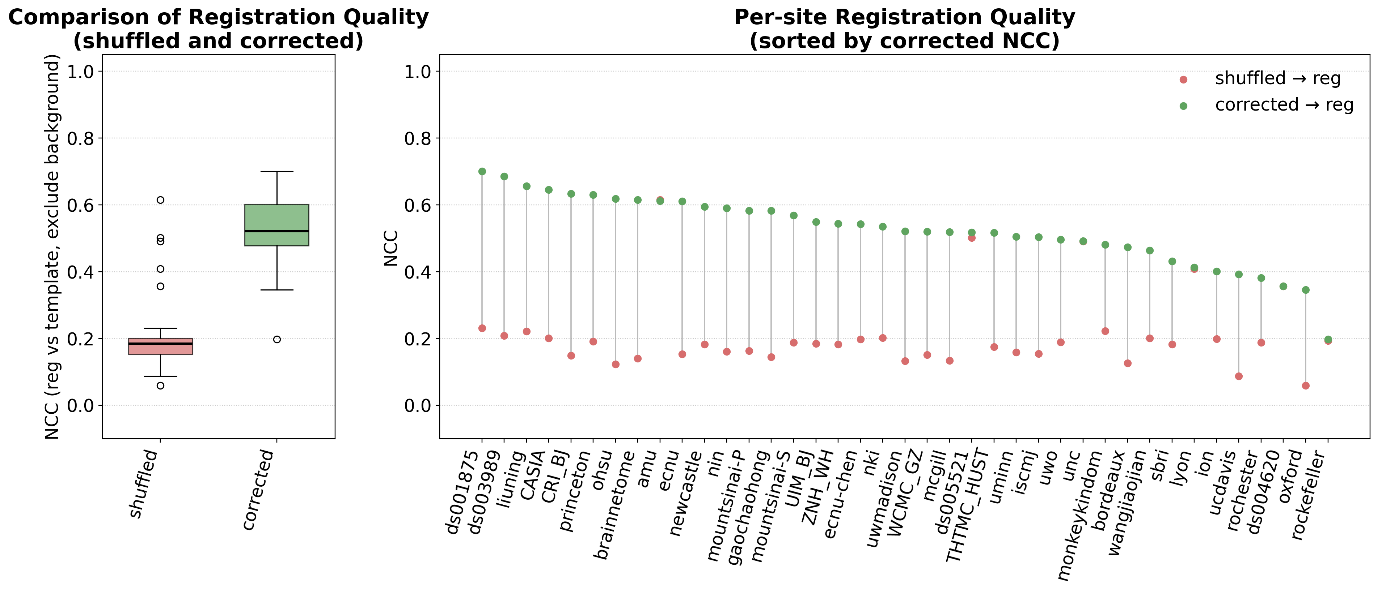

**Supplementary Figure 5. Quantitative evaluation of orientation correction on registration performance.** Left: Mean normalized cross-correlation (NCC) between each image and the template after affine registration, computed within the brain mask, before and after orientation correction (n = 39). Right: Site-wise improvement in NCC after orientation correction, showing consistent gains in registration accuracy across all centers.

#### **Tissue Segmentation**

Model performance was evaluated using 5-fold subject-exclusive cross-validation, where samples from the same subject were strictly prohibited from being split into different folds to avoid data leakage, with an 8:2 training/validation split performed within each acquisition site to maintain site distribution consistency, and the Dice similarity coefficient (DSC) was used as the primary evaluation metric across 18 tissue classes. During inference, we systematically evaluated the effect of different sliding window overlap ratios on segmentation performance under Gaussian-weighted fusion (with Gaussian kernel sigma set to half of the overlap ratio to ensure smooth patch boundary transitions), finding that 60% overlap achieved the best cross-validation average DSC of 0.838, with a 2.95% improvement over the 0% overlap baseline, which was adopted for all subsequent experiments to effectively reduce boundary artifacts while maintaining reasonable inference efficiency. The final model, selected based on the highest average validation DSC across all folds, achieved a mean DSC of 0.838 across all 18 tissue classes on the held-out independent test set, with high performance for large brain structures (cortical gray matter: 0.898, white matter: 0.880) and competitive performance for small subcortical structures (hippocampus: 0.835, amygdala: 0.842, substantia nigra: 0.765); when evaluated on out-of-distribution scans from 4 unseen acquisition sites (OpenNeuro public datasets), the model showed less than 5% DSC drop compared to in-distribution performance, demonstrating strong cross-site generalization ability and robustness to different scanning protocols and hardware platforms.

#### **Registration Benchmarking**

Across all intra-modal experiments, all three tools produced visually acceptable rigid alignment with comparable PSNR and SSIM distributions across parameter configurations. For cross-modal registration, performance varied more substantially: reg_aladin achieved the highest visual pass rate (100%), followed by antsRegistration (97.0%) and FLIRT (93.1%) (Supplementary Fig. 6c).

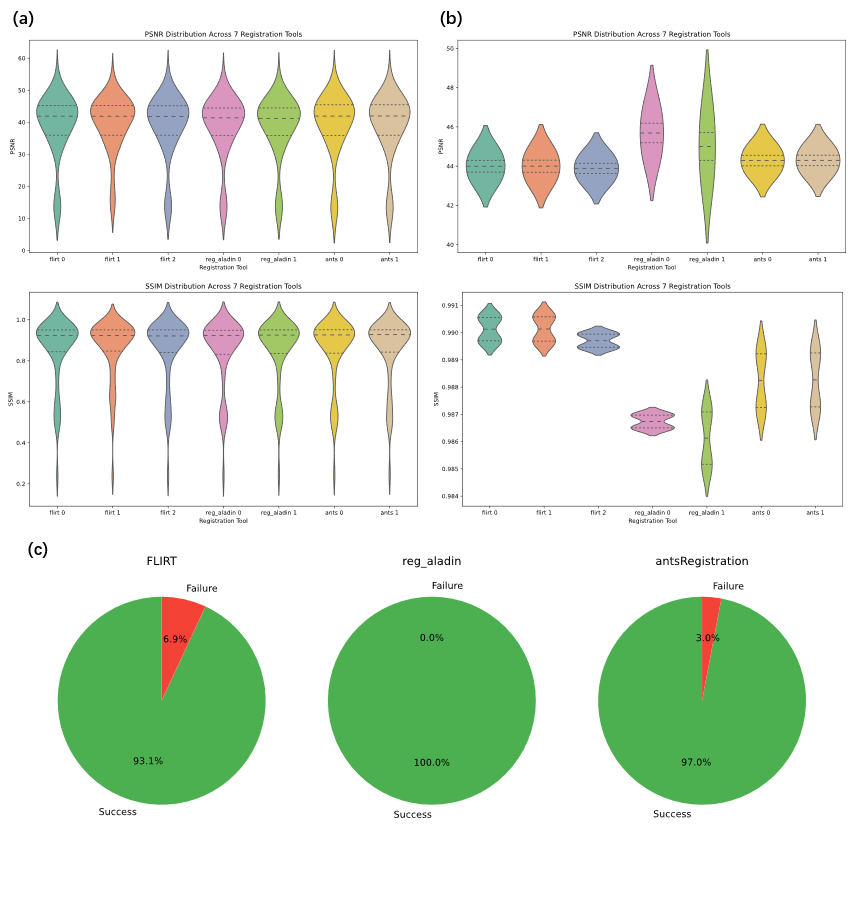

**Supplementary Figure 6. Performance metrics for registration: (a)** PSNR and SSIM for intra-modal T1-to-T1 registration; **(b)** PSNR and SSIM for intra-modal T2-to-T2 registration; **(c)** Visual pass rates of cross-modal (T2-to-T1) registration evaluated for the three tools.

Based on these benchmarking results, we selected FLIRT (param_0, default correlation ratio cost) for all intra-modal rigid alignment steps and NiftyReg reg_aladin (param_1, contrast-enhanced configuration) for cross-modal T2-to-T1 registration in the final MacaSurfer workflow, balancing computational efficiency, robustness, and alignment accuracy across all use cases.

#### **Bias Field Correction Performance**

Here, we demonstrated the impact of the proposed tissue-guided bias field correction method on cortical surface reconstruction, in comparison with standard N4 correction.

The tissue-guided approach better preserved high-frequency intensity variations in deep cortical folds, where standard N4 correction tended to over-smooth intensity profiles. This improvement resulted in more accurate delineation of white matter boundaries, particularly in regions with thin white matter structures.

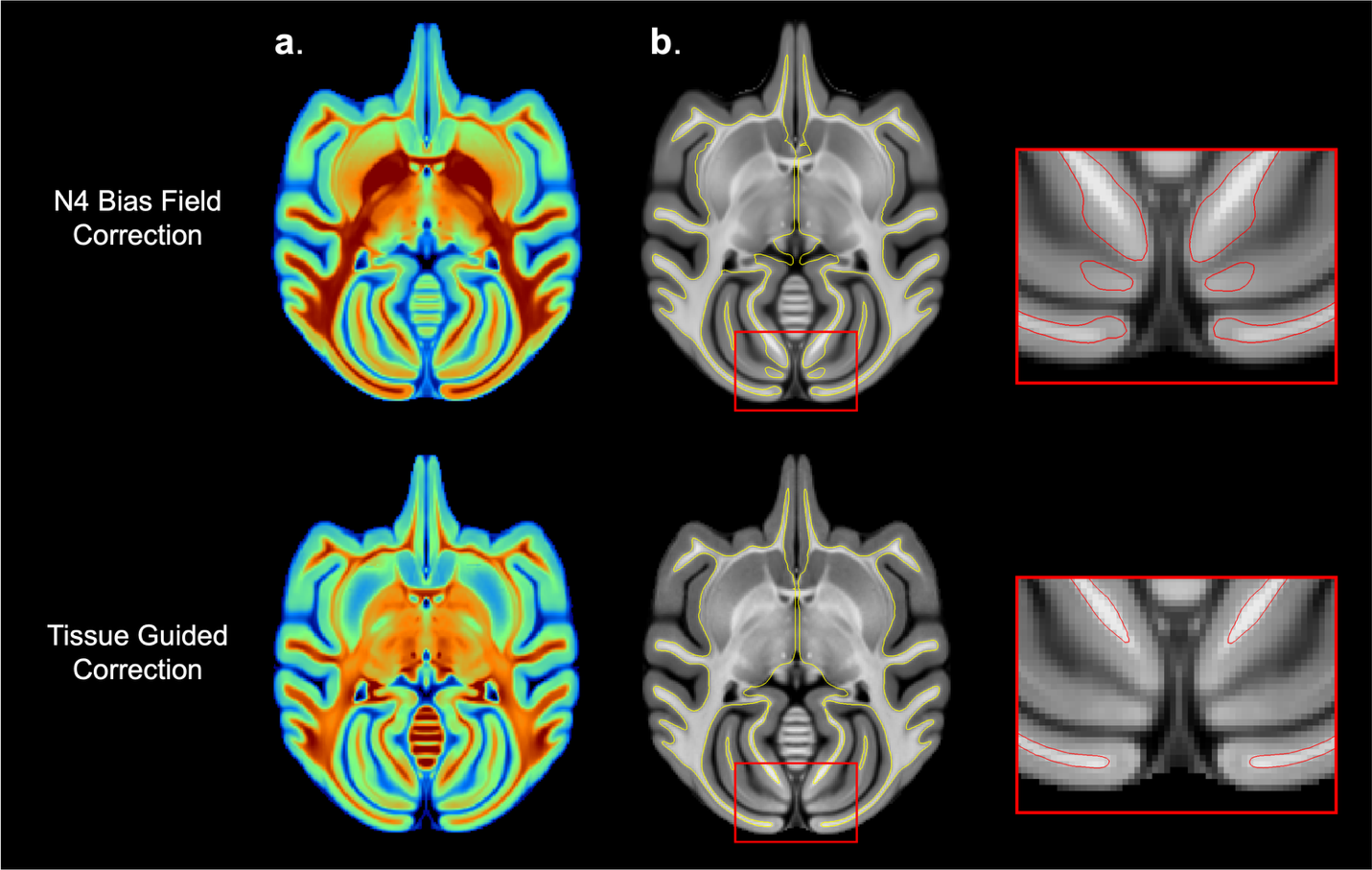

**Supplementary Figure 7. Impact of Tissue-Guided Bias Field Correction on White Surface Generation. (a)** Comparison of bias fields-corrected T1 between standard N4 correction (top) and the proposed Tissue-Guided method (bottom). The proposed method recovers high-frequency variations in deep cortical folds. **(b)** The resulting white matter surface reconstructions. The red insets demonstrate that Tissue-Guided correction prevents boundary erosion in the occipital lobes (bottom right), preserving the continuity of thin white matter strands compared to N4 correction (top right).

#### **White Matter Fixation**

Errors in tissue segmentation—whether from traditional methods or deep learning—can lead to underestimation of white matter, where true white matter regions are partially missed. Such defects introduce topological inconsistencies in the initial white surface, which may be amplified by subsequent automated topology correction procedures.

To address this issue, MacaSurfer incorporates a robust white matter repair strategy based on reverse mapping from the template surface to the subject volume, followed by skeletonization. This approach enables recovery of thin or fragmented white matter structures, providing a more stable initialization for cortical surface reconstruction.

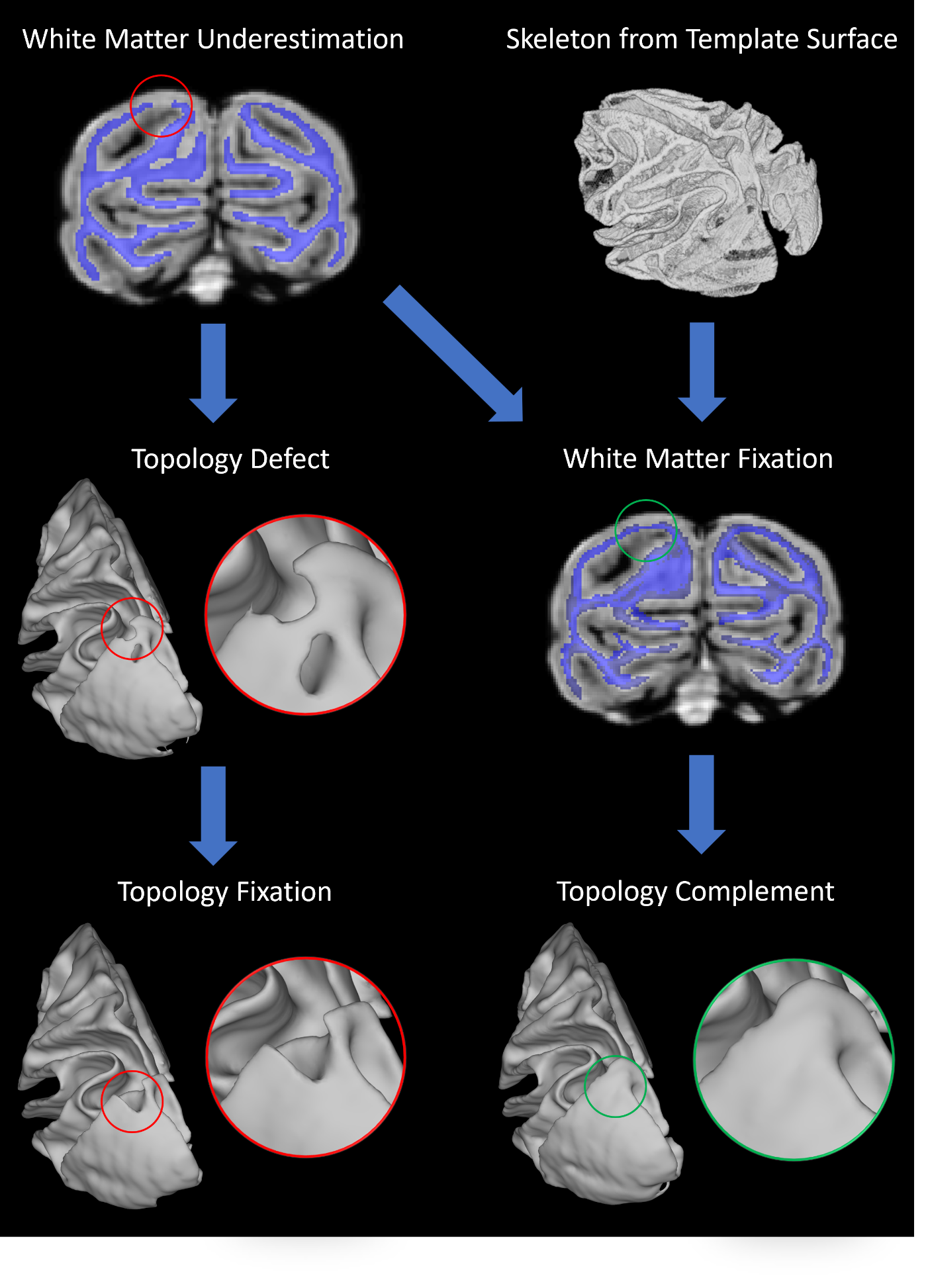

**Supplementary Figure 8. Impact of white matter segmentation errors and correction on white surface reconstruction.** White matter underestimation in tissue segmentation can lead to defects in the reconstructed white surface. For example, cavities in occipital white matter result in corresponding holes in the white surface, while localized missing regions introduce surface indentations. During standard FreeSurfer topology correction, cavities are treated as topological defects, often leading to the removal of adjacent valid connections. Although indentations are not explicitly classified as defects, they can introduce spurious thin connections that mislead topology correction, resulting in the unintended disconnection of anatomically valid structures. MacaSurfer addresses these issues using a template-guided white matter repair strategy. The template-derived surface is mapped to the subject’s volumetric space and skeletonized to reconstruct missing white matter structures. This procedure effectively restores continuity in fragmented regions and resolves holes and indentations in the resulting white surface.

#### **Middle Wall Extraction**

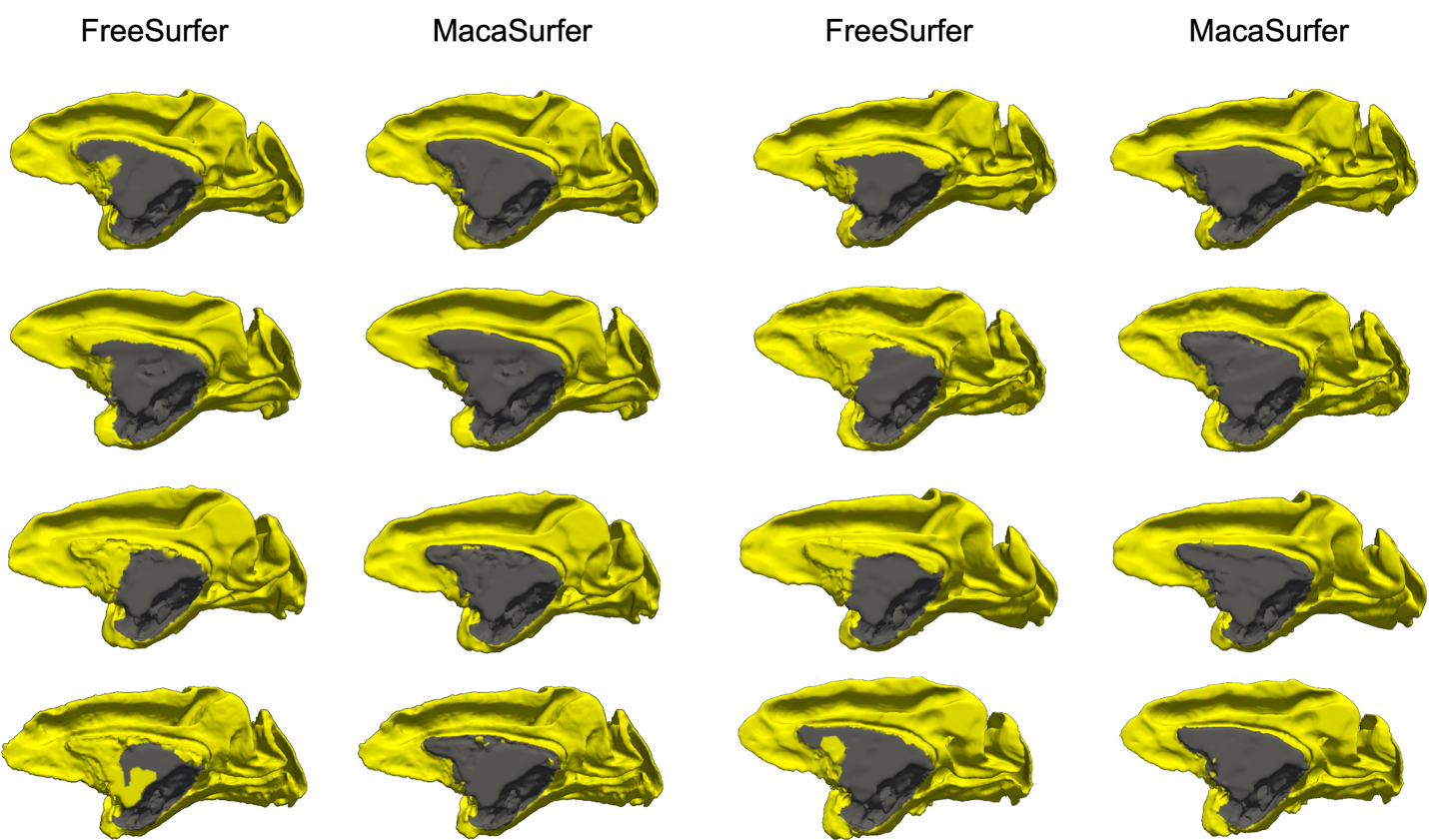

**Supplementary Figure 9. Topological Refinement of the Medial Wall.** Visual comparison of cortical labels derived from standard FreeSurfer (left columns) versus MacaSurfer (right columns). The standard approach frequently misclassifies medial wall structures as cortex. MacaSurfer’s graph-based medial wall extraction explicitly masks these regions (grey), resulting in a topologically correct cortical boundary (yellow) that strictly adheres to anatomical limits.

#### **Surface-Aware Volume Registration**

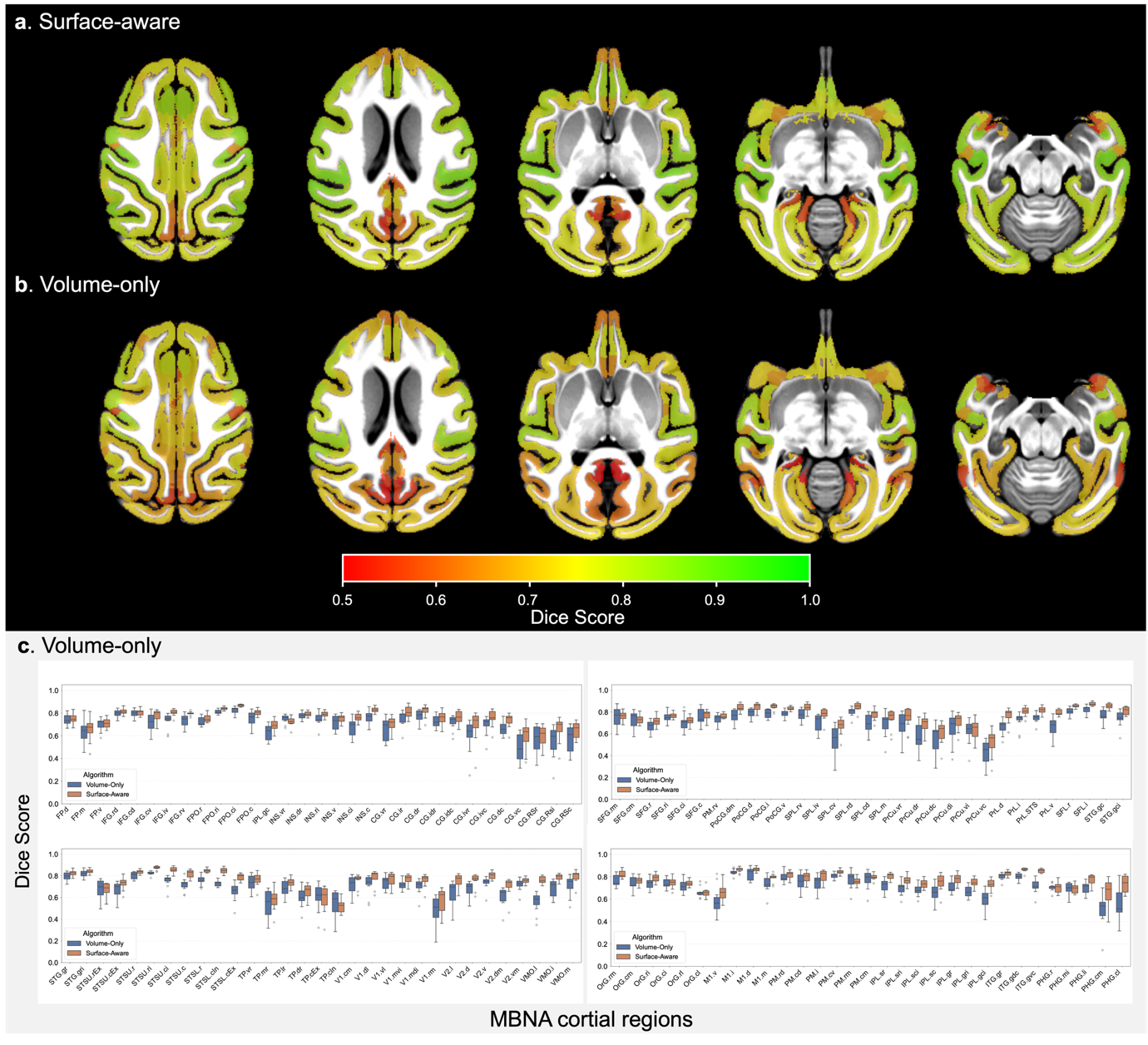

**Supplementary Figure 10. Surface-Aware vs. Volume-Only Registration.** Visual comparison of parcellation alignment with surface-aware (a) and volume-only registration (b). Individual volumetric parcellations were projected to the template space using nonlinear deformation fields estimated by the two registration approaches and visualized in the template volume. Compared with volume-only registration, the surface-aware method shows improved global cortical conformity and better local overlap across most cortical regions, indicating more accurate alignment with cortical anatomy. (c) Boxplots of Dice similarity coefficients for 124 cortical parcels registered to the template. The Surface-Aware strategy (orange) consistently outperforms the Volume-Only approach (blue) across most regions.

### **Supplementary Result 3: Modality Invariance of the MacaSurfer Reconstruction Pipeline**

While multimodal T1-weighted (T1w) and T2-weighted (T2w) integration is widely considered a prerequisite for high-accuracy cortical segmentation, practical acquisition constraints often restrict large-scale cohorts and legacy datasets to T1w-only acquisitions. To evaluate MacaSurfer’s performance under T2w-free conditions, we compared surface reconstructions generated from T1w-only inputs versus combined T1w-T2w inputs in 80 subjects across 12 independent imaging centers.

All statistical comparisons used vertex-wise paired t-tests unless otherwise specified, with false discovery rate (FDR) correction for multiple comparisons (significance threshold p < 0.01, corresponding to -log₁₀(p) > 2). We observed no significant differences in mean cortical thickness across 97.12% of vertices in the left hemisphere and 97.79% of vertices in the right hemisphere. Complementary Kolmogorov-Smirnov (KS) tests confirmed that cortical thickness distributions were statistically indistinguishable for >99.9% of vertices in both hemispheres. Equivalent comparisons for sulcal depth and cortical curvature metrics are provided in Supplementary Table 7. Qualitative validation is shown in Supplementary Fig. 4, which includes direct overlay of T1-only and T1+T2 reconstructed surfaces on native T1w images for visual inspection.

These findings indicate that MacaSurfer’s tissue-guided bias field correction effectively recovers signal in low-contrast regions, rendering the pipeline effectively modality-invariant. This capability allows researchers to derive high-fidelity surface models from historical or simplified acquisition protocols without compromising analytical accuracy.

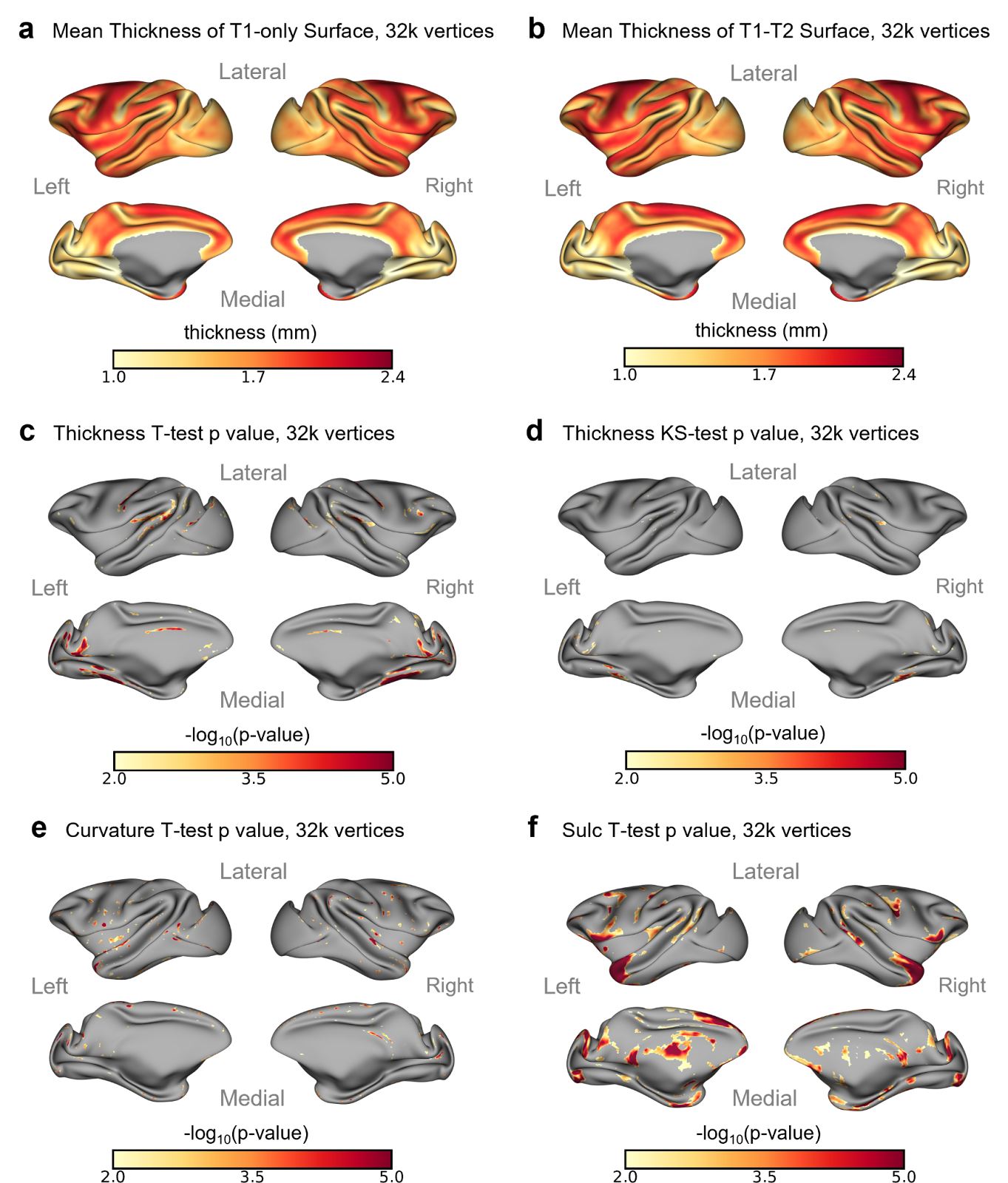

**Supplementary Figure 11. Comparison of T1-only versus multimodal (T1-T2) surface reconstruction. (a)** Mean cortical thickness maps derived solely from T1-weighted images. **(b)** Mean cortical thickness maps derived from combined T1 and T2 images. **(c)** Vertex-wise statistical comparison (paired t-test) showing significant differences in less than 3% of the cortex (colored regions indicate p < 0.01, FDR-corrected), primarily in non-cortical fringe areas. **(d)** Kolmogorov-Smirnov test results demonstrating negligible differences in thickness distributions between the two modalities (significant differences in <0.1% of vertices).

**Supplementary Table 7. Vertex-wise significant difference rates between T1-only and T1+T2 reconstructions**

| **Morphometric metric** | **Hemisphere** | **Significant vertices (paired t-test, FDR-corrected)** | **Significant vertices (KS-test, FDR-corrected)** |
| --- | --- | --- | --- |
| Cortical thickness | Left | 2.88% | 0.09% |
| Cortical thickness | Right | 2.21% | 0.10% |
| Cortical curvature | Left | 1.15% | 0.00% |
| Cortical curvature | Right | 0.55% | 0.00% |
| Sulcal depth | Left | 13.30% | 0.22% |
| Sulcal depth | Right | 11.46% | 0.00% |

*Table note: All tests were performed at the vertex level with FDR correction for multiple comparisons, significance threshold *p* < 0.01 (-log₁₀(*p*) > 2). Values indicate the proportion of vertices showing statistically significant differences between T1-only and combined T1+T2 reconstruction pipelines.*

### **Supplementary Result 4: Visual Report Sample**

We provide an example of the automated visual quality-control report generated by MacaSurfer for a representative non-human primate brain MRI session. The report summarizes the processed dataset at the subject, session, modality, and QC-image levels, and presents all quality-control outputs in an organized, browser-based interface. Users can filter results by subject, session, modality, and QC category, and inspect thumbnail panels for key preprocessing steps, including image orientation, skull stripping, fixed-brain masking, bias-field correction, multimodal registration, template registration and cortical surface reconstruction. This visual report enables rapid review of preprocessing quality across multiple modalities and processing stages, facilitating transparent inspection and efficient identification of potential failure cases before downstream morphometric analyses.

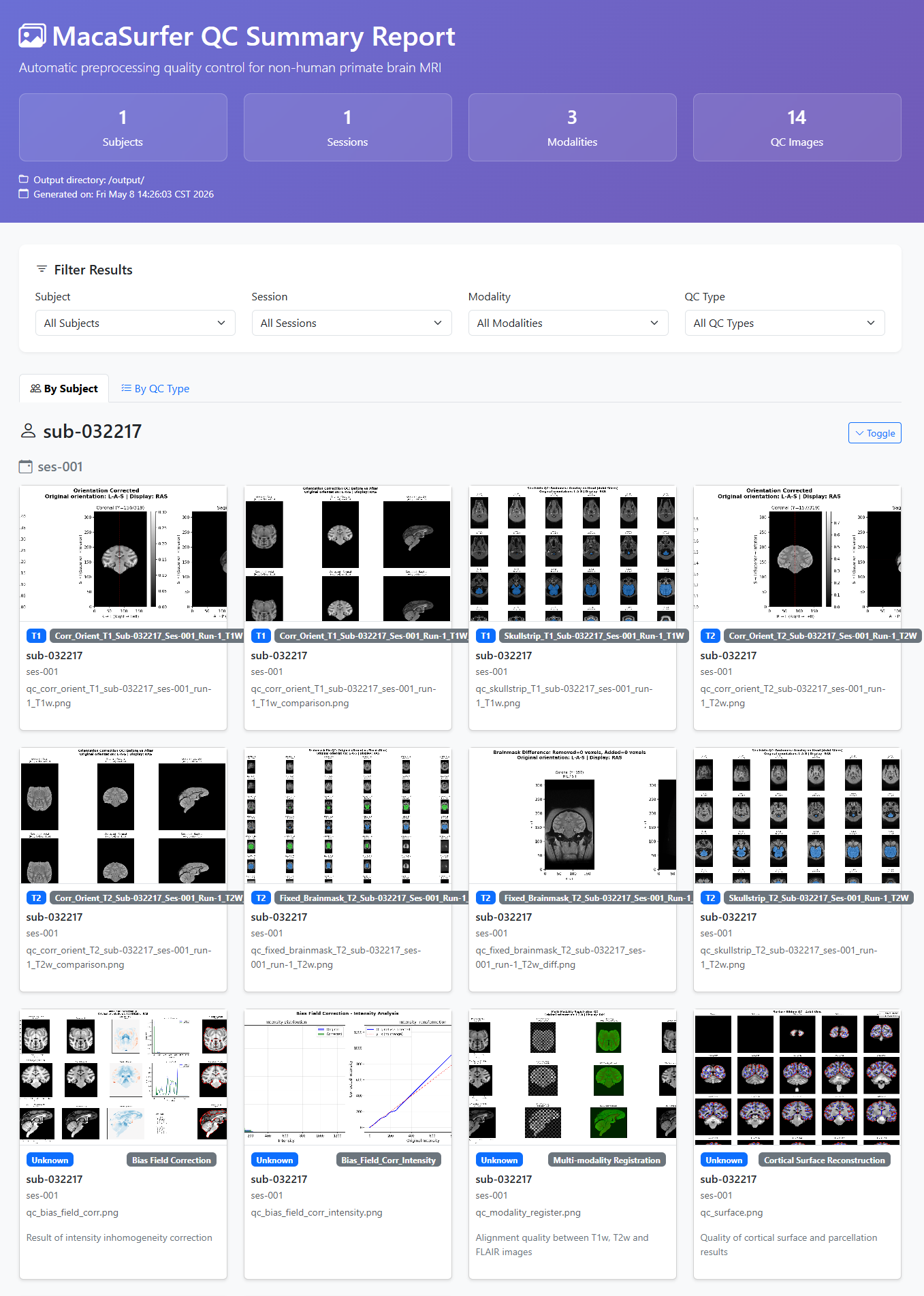

**Supplementary Figure 3. Example visual quality-control report generated by MacaSurfer.** A representative MacaSurfer QC report page showing dataset-level summary statistics, filtering options, and subject/session-specific QC image cards. The displayed outputs include representative checks for orientation correction, skull stripping, brain masking, bias-field correction, multimodal registration and cortical surface reconstruction.

### **Supplementary Dataset Collection**

**INDI PRIME-DE**

We obtained macaque brain imaging data from the PRIMatE Data Exchange (PRIME-DE; <http://fcon_1000.projects.nitrc.org/indi/PRIME/>), a large-scale open-science initiative that aggregates independently acquired non-human primate (NHP) magnetic resonance imaging datasets and openly shares them via the International Neuroimaging Data-sharing Initiative (INDI). PRIME-DE was established to address the challenge that technological and methodological advances in NHP neuroimaging have outpaced data accrual, thereby enabling translational and cross-species comparative neuroscience at scale. The initial release of PRIME-DE consisted of 25 independent data collections aggregated across 22 sites, totaling 217 non-human primates. For the present study, we included data from 23 contributing centers within PRIME-DE, namely: amu, bordeaux, ecnu, ecnu-chen, ion, iscmj, lyon, mcgill, mountsinai-P, mountsinai-S, newcastle, nin, nki, ohsu, oxford, princeton, rochester, rockefeller, sbri, ucdavis, uminn, uwmadison, and uwo. All data collections within PRIME-DE include at least one structural MRI scan and one resting-state functional MRI scan per subject; a subset of collections also provide diffusion MRI and field map data. Detailed scanner specifications, acquisition parameters, and phenotypic information (age, sex, species) for each collection are documented on the PRIME-DE portal.

**UNC-Wisconsin Neurodevelopment Database**

Early postnatal developmental data were obtained from the UNC-Wisconsin Rhesus Macaque Neurodevelopment Database, a publicly available resource characterizing normal postnatal macaque brain development. This longitudinal database includes structural MRI (T1w and T2w) and diffusion tensor imaging (DTI) data acquired from a cohort of 34 typically developing rhesus macaques (mulatta) scanned longitudinally from approximately 2 weeks to 36 months of age. The dataset was generated through a collaboration between the University of North Carolina at Chapel Hill and the University of Wisconsin–Madison, with the goal of providing a normative reference for early brain development in a translational non-human primate model. Detailed descriptions of the study design, animal preparation, and MRI acquisition protocols have been published previously.

**OpenNeuro Datasets**

We additionally curated the following open-access macaque neuroimaging datasets from the OpenNeuro platform ([https://openneuro.org](https://openneuro.org/)), a BRAIN Initiative-designated data archive for the open sharing of human and non-human brain imaging data following FAIR principles:

**ds001875:** A macaque structural and diffusion MRI dataset released under a CC0 license. This dataset was originally generated to construct a macaque connectome for large-scale network simulations using TheVirtualBrain framework and has been used in multiple cross-species comparative neuroimaging studies. The dataset is available at [https://doi.org/10.18112/openneuro.ds001875.v1.0.3](https://doi.org/10.18112/openneuro.ds001875.v1.0.3%5Breference:13%5D%5Breference:14%5D).

**ds003989:** A multimodal MRI dataset including structural, diffusion, and resting-state functional MRI acquired in macaque monkeys. The data were collected at the Centre de Recherche Cerveau et Cognition (CerCo), CNRS, and have been used to investigate the connectivity of the cingulate sulcus visual area (CSv) in macaques (De Castro et al., Cerebral Cortex, 2021). The dataset is available at [https://doi.org/10.18112/openneuro.ds003989.v1.0.0](https://doi.org/10.18112/openneuro.ds003989.v1.0.0%5Breference:17%5D).

**ds004620:** An open-access dataset containing structural MRI data from macaque monkeys, available through the OpenNeuro platform. This dataset has been employed in auditory cortex studies of the macaque, contributing to the characterization of cortical organization in the NHP brain.

**ds005521:** A dataset containing functional and structural MRI data acquired from two alert rhesus macaques (mulatta, 7–8 kg) at the Martinos Imaging Center at Massachusetts General Hospital. Scanning was performed on a 3 Tesla Allegra scanner (Siemens) using a custom-made four-channel send–receive surface coil. The data have been used to investigate color and spatial frequency processing in retinotopic visual areas (V1, V2, V3, V4, MT, and V3a) of the macaque cortex. The dataset is available at <https://openneuro.org/datasets/ds005521>.

**Private Datasets**

In addition to publicly available data, we obtained macaque structural MRI data from six centers at the Chinese Institute for Brain Research, Beijing (namely: CASIA, CRI_BJ, THTMC_HUST, UIM_BJ, WCMC_GZ, and ZNH_WH), and from five collaborating laboratories (Brainnetome, Wangjiaojian, Liuning, and Gaochaohong). The acquisition protocols, sample characteristics, and ethical approvals for these private datasets are detailed separately.

**Supplementary Table 8. Detiled information of all collected sites.**

| **source** | **site** | **# subjects** | **# sessions** | **# T1w** | **# T2w** | **# FLAIR** |
| --- | --- | --- | --- | --- | --- | --- |
| PRIME-DE | amu | 4 | 4 | 4 | 4 | 0 |
|  | bordeaux | 9 | 9 | 9 | 9 | 0 |
|  | ecnu | 4 | 4 | 4 | 1 | 0 |
|  | ecnu-chen | 10 | 10 | 11 | 0 | 0 |
|  | ion | 8 | 9 | 9 | 0 | 0 |
|  | iscmj | 4 | 14 | 14 | 0 | 0 |
|  | lyon | 4 | 4 | 9 | 0 | 0 |
|  | mcgill | 3 | 8 | 21 | 1 | 0 |
|  | mountsinai-P | 9 | 9 | 27 | 5 | 0 |
|  | mountsinai-S | 5 | 5 | 50 | 5 | 0 |
|  | newcastle | 14 | 22 | 22 | 10 | 0 |
|  | nin | 2 | 29 | 66 | 0 | 0 |
|  | nki | 2 | 36 | 81 | 0 | 0 |
|  | ohsu | 2 | 16 | 16 | 16 | 0 |
|  | oxford | 20 | 20 | 92 | 0 | 0 |
|  | princeton | 2 | 8 | 12 | 2 | 0 |
|  | rochester | 3 | 3 | 3 | 0 | 0 |
|  | rockefeller | 6 | 6 | 6 | 0 | 0 |
|  | sbri | 22 | 22 | 47 | 3 | 0 |
|  | ucdavis | 19 | 19 | 19 | 19 | 0 |
|  | uminn | 2 | 2 | 2 | 2 | 0 |
|  | unc | 34 | 161 | 163 | 162 | 0 |
|  | uwmadison | 584 | 584 | 584 | 0 | 0 |
|  | uwo | 12 | 12 | 12 | 12 | 0 |
| OpenNeuro | ds001875 | 9 | 9 | 9 | 0 | 0 |
|  | ds003989 | 3 | 3 | 13 | 0 | 0 |
|  | ds004620 | 8 | 8 | 8 | 0 | 0 |
|  | ds005521 | 2 | 2 | 2 | 0 | 0 |
| Collaborating Laboratories | wangjiaojian | 24 | 24 | 24 | 0 | 0 |
|  | brainnetome | 1 | 11 | 11 | 11 | 0 |
|  | gaochaohong | 10 | 30 | 30 | 0 | 0 |
|  | liuning | 24 | 24 | 24 | 24 | 0 |
|  | monkeykindom | 12 | 44 | 44 | 1 | 0 |
| Chinese Institute for Brain Research | CRI_BJ | 32 | 33 | 50 | 37 | 4 |
|  | UIM_BJ | 12 | 72 | 58 | 59 | 0 |
|  | THTMC_HUST | 1 | 2 | 6 | 6 | 0 |
|  | CASIA | 4 | 4 | 8 | 6 | 4 |
|  | WCMC_GZ | 5 | 5 | 5 | 0 | 9 |
|  | ZNH_WH | 35 | 85 | 261 | 135 | 16 |
| **Total** | **39 sites** | **966** | **1372** | **1836** | **530** | **33** |

### **Supplementary Normative Modeling Data**

Normative models were constructed from a multi-site cohort of 835 neurologically healthy macaques (1,145 imaging sessions, 26 sites), predominantly rhesus macaques (Macaca mulatta; n = 809) with a smaller subset of cynomolgus macaques (Macaca fascicularis; n = 26). Ages ranged from birth to 23 years. The largest site-specific contributions were from the University of Wisconsin–Madison (n = 583), University of North Carolina at Chapel Hill (n = 158), and ZNH Wuhan (private site, n = 73). Animals with a history of neurological disease, traumatic brain injury, developmental abnormality, or visible MRI pathology were excluded. Complete site-level demographic details are provided in Supplementary Table 9.

**Supplementary Table 9. Detailed information of all sites for normative modeling.**

| **site** | **# subjects** | **# sessions** | **age**  **min** | **age**  **max** | **age**  **mean** | **sex** | **breed** |
| --- | --- | --- | --- | --- | --- | --- | --- |
| amu | 4 | 4 | 7 | 8 | 7.5 | F=1, M=3 | mulatta=4 |
| bordeaux | 9 | 9 | 3 | 23 | 10 | F=4, M=5 | fascicularis=9 |
| ecnu | 4 | 4 | 2.67 | 3.75 | 3.4 | M=4 | mulatta=4 |
| ion | 8 | 9 | 3.8 | 5.81 | 5.11 | F=1, M=8 | fascicularis=4, mulatta=5 |
| liuning | 24 | 24 | 9 | 15 | 11.92 | F=24 | mulatta=24 |
| mcgill | 3 | 8 | 10 | 12 | 10.25 | F=8 | fascicularis=7, mulatta=1 |
| mountsinai-P | 9 | 9 | 3.4 | 8 | 4.8 | F=1, M=8 | fascicularis=1, mulatta=8 |
| mountsinai-S | 5 | 5 | 5.3 | 6.3 | 5.94 | M=5 | mulatta=5 |
| newcastle | 14 | 22 | 3.9 | 13.14 | 8 | F=2, M=20 | Mulatta=22 |
| nin | 2 | 29 | 5 | 6.5 | 6.03 | M=29 | mulatta=29 |
| nki | 2 | 36 | 6 | 7 | 6.75 | F=27, M=9 | mulatta=36 |
| ohsu | 2 | 16 | 5 | 5 | 5 | M=16 | mulatta=16 |
| oxford | 20 | 20 | 2.41 | 6.72 | 4.01 | M=20 | mulatta=20 |
| princeton | 2 | 8 | 3 | 3 | 3 | M=8 | mulatta=8 |
| rochester | 3 | 3 | 3 | 3 | 3 | F=1, M=2 | fascicularis=3 |
| rockefeller | 6 | 6 | 4 | 4 | 4 | M=6 | fascicularis=1, mulatta=5 |
| sbri | 22 | 22 | 3.6 | 13.9 | 8.4 | F=13, M=9 | fascicularis=6, mulatta=16 |
| THTMC_HUST | 1 | 2 | 8.95 | 10.47 | 9.71 | M=2 | mulatta=2 |
| ucdavis | 19 | 19 | 18.6 | 22.5 | 20.38 | F=19 | mulatta=19 |
| UIM_BJ | 12 | 55 | 8.21 | 13.31 | 10.67 | M=55 | mulatta=55 |
| uminn | 2 | 2 | 10 | 10 | 10 | F=2 | mulatta=2 |
| unc | 33 | 160 | 0.0425 | 3 | 1.05 | F=75, M=85 | mulatta=160 |
| uwmadison | 583 | 583 | 0.835616438 | 4.42 | 1.91 | F=262, M=321 | mulatta=583 |
| uwo | 12 | 12 | 4 | 8 | 5.33 | M=12 | mulatta=12 |
| WCMC_GZ | 5 | 5 | 8.07 | 8.83 | 8.65 | M=5 | mulatta=5 |
| ZNH_WH | 33 | 72 | 4.52 | 21.39 | 10.53 | M=72 | mulatta=72 |
| **26 sites** | **839** | **1144** | **0.0425** | **23** |  |  |  |

### **Supplementary Evaluation Data**

To systematically verify the stability, accuracy, and generalizability of the MacaSurfer pipeline and its standardized outputs, we conducted a series of targeted validation experiments across core functional modules. The datasets, selection criteria, and experimental designs employed for these quantitative and qualitative evaluations are detailed below.

**Deep learning segmentation training:** For the development of the deep learning segmentation model, surface reconstruction was performed for every session of each subject. In cases where a session contained multiple runs, a single label was shared across runs since they were already aligned via rigid-body registration. The training set comprised 2,157 scans from 39 acquisition centers, with 1,699 T1-weighted, 425 T2-weighted and 33 FLAIR scans. Four public datasets from OpenNeuro were reserved as an independent cross-site test set. Detailed dataset composition and site information are provided in Supplementary Table 10.

**Supplementary Table 9. Detailed information of all sites for tissue segmentation model.**

| **usage** | **site** | **# subjects** | **# T1w** | **# T2w** | **# FLAIR** |
| --- | --- | --- | --- | --- | --- |
| Training / Validation | amu | 4 | 4 | 4 | 0 |
|  | bordeaux | 9 | 9 | 9 | 0 |
|  | brainnetome | 1 | 11 | 11 | 0 |
|  | CASIA | 4 | 8 | 3 | 2 |
|  | CRI_BJ | 32 | 50 (4 excluded) | 22 (1 excluded) | 11 |
|  | ecnu | 4 | 4 | 1 | 0 |
|  | ecnu-chen | 10 | 11 | 0 | 0 |
|  | gaochaohong | 10 | 28 | 0 | 0 |
|  | ion | 8 | 9 | 0 | 0 |
|  | iscmj | 4 | 14 (8 excluded) | 0 | 0 |
|  | liuning | 24 | 24 | 24 | 0 |
|  | lyon | 4 | 9 | 0 | 0 |
|  | mcgill | 3 | 21 (1 excluded) | 1 | 0 |
|  | monkeykindom | 12 | 44 | 1 | 0 |
|  | mountsinai-P | 9 | 27 | 5 | 0 |
|  | mountsinai-S | 5 | 50 (3 excluded) | 5 | 0 |
|  | newcastle | 14 | 22 | 10 | 0 |
|  | nin | 2 | 66 | 0 | 0 |
|  | nki | 2 | 81 | 0 | 0 |
|  | ohsu | 2 | 16 | 16 | 0 |
|  | oxford | 20 | 92 | 0 | 0 |
|  | princeton | 2 | 12 | 2 | 0 |
|  | rochester | 3 | 3 | 0 | 0 |
|  | rockefeller | 6 | 6 | 0 | 0 |
|  | sbri | 22 | 47 | 3 | 0 |
|  | THTMC_HUST | 1 | 6 | 2 | 0 |
|  | ucdavis | 19 | 19 | 19 | 0 |
|  | UIM_BJ | 12 | 58 (4 excluded) | 55 (1 excluded) | 1 |
|  | uminn | 2 | 2 | 2 | 0 |
|  | unc | 33 | 162 | 157 (1 excluded) | 0 |
|  | uwmadison | 583 | 583 | 0 | 0 |
|  | uwo | 12 | 12 | 12 | 0 |
|  | wangjiaojian | 24 | 24 | 0 | 0 |
|  | WCMC_GZ | 5 | 5 | 0 | 5 |
|  | ZNH_WH | 36 | 237 (89 excluded) | 67 (3 excluded) | 14 |
| OOD  Test | ds001875 | 9 | 9 | 0 | 0 |
|  | ds003989 | 3 | 13 | 0 | 0 |
|  | ds004620 | 8 | 8 | 0 | 0 |
|  | ds005521 | 2 | 2 | 0 | 0 |
| **Total** | **39 sites** | **947** | **1808 (109 excluded)** | **431 (6 excluded)** | **33** |

**Robustness assessment via noise injection:** Experiments for robustness evaluation were conducted using raw data from subject "Jekyll" from the Site-Monkey Kingdom center. This specific dataset was selected for its exceptionally high raw image quality, which allowed for controlled noise addition without interference from prior correction artifacts.

**Intra-subject reproducibility:** Reproducibility was evaluated using data from two subjects at the Site-NIN center, selected for having the highest number of repeat scans. To ensure an unbiased assessment, multiple runs from the same subject were reconstructed independently rather than being pooled.

**Tissue-guided bias field correction:** Qualitative visualization of bias field correction was performed using the MEBRAINS template. This template was chosen specifically for its extreme B1-field inhomogeneity in the occipital lobe—a significant challenge that conventional low-frequency bias field estimation algorithms typically fail to resolve.

**Surface-aware volumetric registration:** Data from Site-MonkeyKingdom were utilized for surface-aware volumetric registration. This site provided superior image contrast and the most accurate manually vetted surface reconstructions. As our registration framework integrates both volumetric and surface-based information, this dataset provides an optimal benchmark for evaluating the algorithm's performance.
